# The ethanol tolerance in *Saccharomyces cerevisiae* under a phenomics perspective

**DOI:** 10.1101/2021.08.04.455136

**Authors:** Ivan Rodrigo Wolf, Lucas Farinazzo Marques, Lauana Fogaça de Almeida, Lucas Cardoso Lázari, Leonardo Nazário de Moraes, Luiz Henrique Cardoso, Camila Cristina de Oliveira Alves, Rafael Takahiro Nakajima, Amanda Piveta Schnepper, Marjorie de Assis Golim, Thais Regiani Cataldi, Jeroen G. Nijland, Camila Moreira Pinto, Matheus Naia Fioretto, Rodrigo Oliveira Almeida, Arnold J. M. Driessen, Rafael Plana Simōes, Mônica Veneziano Labate, Rejane Maria Tommasini Grotto, Carlos Alberto Labate, Ary Fernandes Junior, Luís Antonio Justulin, Rafael Luiz Buogo Coan, Érica Ramos, Fabiana Barcelos Furtado, Cesar Martins, Guilherme Targino Valente

**Affiliations:** Department of Bioprocess and Biotechnology. School of Agriculture. São Paulo State University (UNESP). Botucatu, SP. Brazil; Department of Structural and Functional Biology, Institute of Biosciences. São Paulo State University (UNESP). Botucatu, SP. Brazil; Laboratory of Applied Biotechnology, São Paulo State University (UNESP), Clinical Hospital of the Medical School, Botucatu, Brazil; Biomedical Sciences Institute, Department of Parasitology. University of São Paulo (USP). São Paulo, SP, Brazil; Laboratório Max Feffer de Genética de Plantas, Escola Superior de Agricultura Luiz de Queiroz, Universidade de São Paulo (USP). Piracicaba, SP, Brazil; Molecular Microbiology, Groningen Biomolecular Sciences and Biotechnology Institute and Zernike Institute for Advanced Materials, University of Groningen, Nijenborgh 7, 9747 AG Groningen, The Netherlands; Instituto Federal de Educação, Ciência e Tecnologia do Sudeste de Minas Gerais - Campus Muriaé. Muriaé, MG, Brazil; Laboratory of Bacteriology, Department of Chemical and Biological Sciences, São Paulo State University (Unesp), Institute of Biosciences, Botucatu; Department of Biophysics and Pharmacology. Institute of Biosciences. São Paulo State University (UNESP). Botucatu, SP, Brazil; Max Planck Institute for Heart and Lung Research. Bad Nauheim, Hessen. Germany

**Keywords:** OMICs, data integration, systems biology, lncRNAs, lncRNA-protein interactions, membraneless, CRISPR-Cas9

## Abstract

Ethanol (EtOH) is a substantial stressor for *Saccharomyces cerevisiae*. Data integration from strains with different phenotypes, including EtOH stress-responsive lncRNAs, are still not available. We covered these issues seeking systems modifications that drive the divergences between higher (HT) and lower (LT) EtOH tolerant strains under their highest stress conditions. We showed that these phenotypes are neither related to high viability nor faster population rebound after stress relief. LncRNAs work on many stress-responsive systems in a strain-specific manner promoting the EtOH tolerance. Cells use membraneless RNA/protein storage and degradation systems to endure the stress harming, and lncRNAs jointly promote EtOH tolerance. CTA1 and longevity are primer systems promoting phenotype-specific gene expression. The lower cell viability and growth under stress is a byproduct of sphingolipids and inositol phosphorylceramide dampening, acerbated in HTs by sphinganine, ERG9, and squalene overloads; LTs diminish this harm by accumulating inositol 1-phosphate. The diauxic shift drives an EtOH buffering by promoting an energy burst under stress, mainly in HTs. Analysis of mutants showed genes and lncRNAs in three strains critical for their EtOH tolerance. Finally, longevity, peroxisome, energy and lipid metabolisms, RNA/protein degradation and storage systems are the main pathways driving the EtOH tolerance phenotypes.

## Introduction

*Saccharomyces cerevisiae* is the main organism used in ethanol (EtOH) production due to its fast growth and high resistance to stressful environments. However, alcoholic fermentation by-products negatively impact cells, such as EtOH, oxidative stresses, and others (Auesukaree 2017).

EtOH yield affects the physiological condition of yeast, altering the growth rate and cell survival (Stanley et al. 2010a). EtOH stress alters the plasma membrane fluidity and the permeability for ions, harming intracellular homeostasis and acidifies the cells. That stress also results in protein dysfunction, facilitates nucleotides, amino acids, and potassium efflux and reduces glucose, maltose, ammonia, and amino acids intake (Ma and Liu 2010). EtOH-stressed cells activate response mechanisms modulating multiple biological processes to avoid those side effects (Yang and Tavazoie 2020).

Massive information from OMICs, and network analysis revealed that the EtOH stress response in yeast is a complex mechanism involving several pathways. For instance, the transcriptome, proteome, and metabolome landscapes revealed imbalances of many processes associated with stress responses (homeostasis, heat shock proteins, and redox balance), cell structures (plasma membrane components, and vacuolar functions), metabolisms (oxidative stress, amino acids, energy, flavoproteins, carbon, volatile compounds, polyphenols, TCA cycle, and L-aspartate), signal transduction, and RNA and protein synthesis (Stanley et al. 2010b; Lourenço et al. 2013; Lewis et al. 2014; Santos et al. 2017; Cheng et al. 2019; Li et al. 2019a). Disrupting regulatory network by mutating stress response transcription factors showed that glycolysis is related to EtOH tolerance (Liang et al. 2021). Moreover, combining transcriptome, network, and cell morphology data of one strain under EtOH stress revealed that the amino acids, trehalose, ergosterol, thiamine metabolisms, and the ubiquitin-proteasome proteolytic pathway are essential for that stress response (Li et al. 2017). Proteomics, genomic sequencing, and network analysis of evolved strains under EtOH stress showed modifications on many metabolic pathways, including energy and lipid metabolism (Šoštarić et al. 2021). Analysis of protein-protein interactions and regulatory networks showed network changings under EtOH stress, embracing many genes related to lipid metabolism, transport, respiration, stress response, and nutrient-sensing mechanisms (Kasavi et al. 2014, 2016). Remarkably, the EtOH stress response seems to be strain-specific, add-on complexity to understand this phenomenon. For instance, EtOH prompts a high diversity of network community organization, distinct metabolic adaptabilities, and different stress responses according to polymorphisms and chromosomal rearrangements (Kang et al. 2019).

Approximately 85% of *S. cerevisiae* genome is expressed under basal conditions, including a bulk of long non-coding RNA (lncRNAs) (David et al. 2006). LncRNAs play a role in promptly responding to external stimuli (Yamashita et al. 2016), mainly working on gene expression regulation and metabolism (Yamashita et al. 2016; Novačić et al. 2020; Balarezo-Cisneros et al. 2021).

LncRNAs may bind to DNA, RNAs, or proteins (Novačić et al. 2020). The lncRNA-protein interactions foster the framework of macromolecular complex assembly, recruitment of transcription factors, guiding chromatin modifiers, or performing a competitive binding against DNA-binding proteins (Kino et al. 2010; Kornienko et al. 2013; Ferrè et al. 2016; Li et al. 2019b). Yeast lncRNAs can activate or block gene expression in either *cis* or *trans* mode by acting on histone or scaffolding RNPs (Niederer et al. 2017; Till et al. 2018; Novačić et al. 2020). Additionally, lncRNA may act as bait blocking the target-protein function or as scaffolders, assisting in protein complex organization either positively or negatively (Wang and Chang 2011).

We found EtOH stress-responsive lncRNAs of 6 yeast strains with two phenotypes (a higher and a lower EtOH tolerant strains). By binding to several proteins, these lncRNAs seem to work on the cell wall, cell cycle, growth, longevity, surveillance, ribosome biogenesis, transcription, among others, in a strain-specific manner (Marques et al. 2021). Cell cycle dynamic modeling and gene inactivation experiments performed by us revealed that lncRNAs transcr_9136 and transcr_10883 are responsible for skipping the cell cycle arrest under EtOH stress (Lázari et al. 2021).

The goal of systemic analysis is to fully view multifactorial characteristics of complex systems. Despite the progress concerning EtOH tolerance in yeast, massive data integration from strains with different phenotypes is still not available. Moreover, there is neither report investigating the role of EtOH stress-responsive lncRNAs, nor their interactions. Therefore, herein we focus on EtOH tolerance integrating data from OMICs, cell and molecular biology, bioinformatics, modeling, networks and mutant analysis seeking features that could differentiate the higher and lower EtOH tolerant phenotypes.

Experiments using 3 higher (HT: BMA64-1A, BY4742, and X2180-1A) and 3 lower (LT: SEY6210, BY4741, and S288C) EtOH tolerant strains showed that these phenotypes are neither related to the high cell viability nor to the faster population rebound after stress-relief. A combination of pathways is affected by the EtOH stress, which longevity, peroxisome, RNA biology, energy and lipid metabolisms, RNA/protein degradation and storage mechanisms as the main ones to drive the EtOH tolerance phenotypes. In many cases, lncRNAs work on these systems in a strain-specific manner: *e.g*., the transcr_20548-Nrp1p interaction in BMA64-1A is related to its lower rebound after stress; the action of transcr_18666 on trehalose in S288C leads to its higher rebound and viability. Cells under stress use membraneless and RNA/protein storage and degradation systems, jointly acting with lncRNAs to endure stress harming promoting EtOH tolerance. Only basal pathways are kept active under stress to prepare cells for stress relief. Interestingly, CTA1 flows signals to the longevity path, which are the first to exhibit expression phenotype divergences. The lower cell viability and growth under stress may be a by-product of sphingolipids and inositol phosphorylceramide synthesis reduction, acerbated by sphinganine, ERG9, and squalene overload in HTs. Conversely, LTs diminish this harmful effect by accumulating inositol 1-phosphate. We draw the EtOH buffering model: the diauxic shift mechanism promote an energy burst under stress, which HTs seems to improve this boost by reducing the oxaloacetate and enhancing SDH activity. Finally, analysis of mutants showed that CTA1, IXR1, lncRNAs transcr_20548, transcr_6448 (BMA64-1A), transcr_3536 (SEY6210), and transcr_10027 (BY4742) are critical for their EtOH tolerance.

## Material and Methods

It was defined the highest EtOH level supported for 13 different *S. cerevisiae* strains obtained from Euroscarf (European Saccharomyces cerevisiae Archive for Functional Analysis) and NRBP (National Bioresources Project). Selected strains were classified into higher tolerant (HT) or lower EtOH tolerant (LT) phenotypes (**Figure 1A-C**). Time-course experiment (1 h, 2h, and 4h) was performed for BMA64-1A (HT) and S288C (LT) under their highest EtOH tolerance conditions (**Figure 1B**). Further, experiments were scaled accordingly (**Figure 1D**).

**Figure 1:**
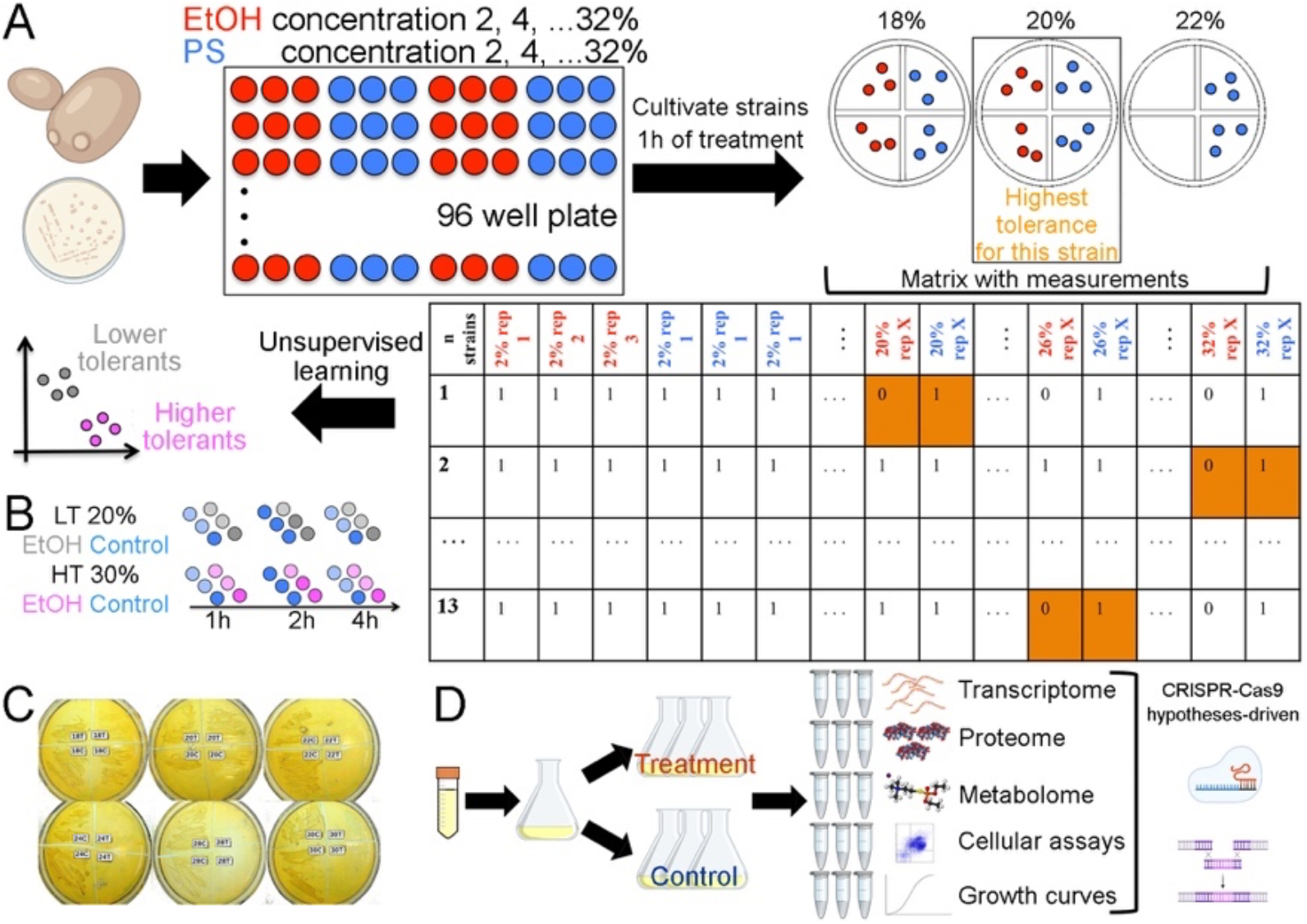
Experimental design. **A:** setting the highest EtOH tolerance level and phenotypes. **PS:** physiological solution; **Red and blue circles:** treatment and control, respectively. **Orange box:** example of the highest EtOH level; **B:** time-course. **Light circles:** selected samples for RNA-Seq; **C:** plates to exemplify to determine the highest EtOH tolerance (S288C); **C and T:** control and treatment, respectively; **D:** further experiments and hypothesis testing using mutants.

## Results

### 1. Cell growth analysis and defining the ethanol tolerances

The highest EtOH tolerance level supported by each strain ranged from 18% to 30% EtOH. The BMA64-1A, BY4742, and X2180-1A exhibits the HT phenotype, whereas SEY6210, BY4741, and S288C composed the LTs (**Table 1**). There was a negative correlation between the EtOH tolerance level and the population rebound after the stress relief. For instance, S288C has the fastest rebound, while the BMA64-1A presents an inverse profile (**Figure 2A, and C**).

**Table 1:** Strains description, cell viability, and SDH assays. **1:** genome sequenced; **2:** used in the timecourse experiment; *: statistical significant data (p-value <0.05).

| Strain or group | EtOH tol. (%) | Phenotype | SDH p-value | Cell viability p-value |
| --- | --- | --- | --- | --- |
| BMA64-1A<sup>1,2</sup> | 30 | HT | 0.002* | 3.9E-13* |
| BY4742 | 26 | HT | 0.001* | 2.2E-16* |
| X2180-1A | 24 | HT | 0.205 | 2.2E-16* |
| BY4741 | 22 | LT | 0.496 | 0.8554 |
| SEY6210 | 20 | LT | 0.718 | 2.6E-07* |
| S288C<sup>2</sup> | 20 | LT | 0.097 | 0.0015* |
| All HTs | - | - | 0.025* | 2.2E-16* |
| All LTs | - | - | 0.135 | 0.0195* |

**Table 2:** Pearson correlation between relative pH and K within different EtOH stresses. The p is out of parenthesis (p-value). Low, mid and high EtOH levels are 5%, 10% and 15% for S288C and 8%, 16% and 23% for BMA64-1A (**Figure 2F**).

| Strain | Low EtOH level | Mid EtOH level | High EtOH level |
| --- | --- | --- | --- |
| BMA64-1A | -0.961 (<0.001) | -0.873 (<0.0001) | -0.799 (<0.01) |
| S288C | -0.968 (<0.0001) | -0.988 (<0.005) | -0.810 (<0.01) |

**Figure 2:**
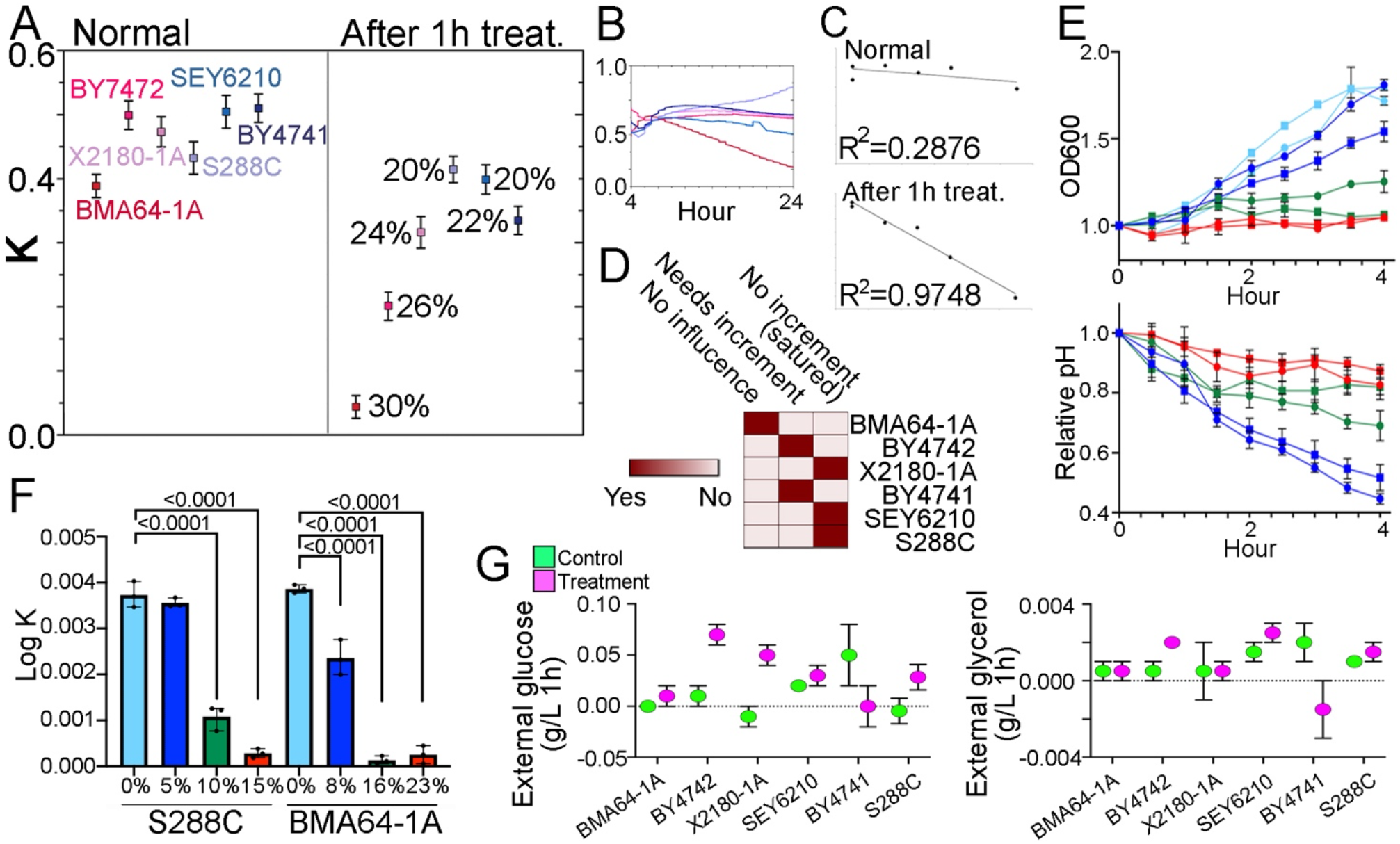
Growth analysis. **A:** population growth rate in YPD without any stressor (left plot) and after the highest EtOH stress treatment for 1h (right plot) (the stress-relief). The percentages are the highest EtOH level tested; **B:** the growth curve along 24h under the highest EtOH level; **C:** regression for the highest EtOH level and K reported on “A”; **D:** overview concerning impact of spermidine in growth summarizing the growth reported with and without spermidine (**Supplementary Figure 6D-E**) and spermidine abundances from metabolomics (**Supplementary Data 8**); **E-F:** the relationship between cell growth, EtOH stress levels, and the medium acidification. Realize that the pH and OD_600_ are corrected and normalized rather than raw data (details in **equations 1** and **2** in **Supplementary Text**). p-value adjusted is shown on bars. The “E” colors match the “F”; **G:** glucose and glycerol measurements. The Y-axis is the glucose consumption and glycerol production rate estimated after 1 h.

There was a negative correlation between the relative pH and K parameters (growth) within each EtOH level (**Table 1**). The lower EtOH stresses had a small impact on BMA64-1A and S288C growth and, hence, higher medium acidification. The high and mid EtOH stress prompts a faster steady-state on cell growth and medium acidification, suggesting that the cytosol may be acidifying under these conditions. Altogether, the medium acidification stops as the population does not increase under mid and high EtOH stress (**Figure 2E-F**). EtOH hampers the control of cytosol’s pH in yeast. Then, cells grown in glucose quickly acidify the medium, and the cytosol acidifies as cells reach stationary-phase or under glucose starvation conditions (Kane 2016; Kuroda et al. 2019). The EtOH stress decreased the glucose intake for almost all strains, and only BMA64-1A, X2180-1A, and BY4741 presented no shift or reduced external glycerol yield (**Figure 2G**).

Spermidine is annotated as part of four KEGG enriched pathways (**Supplementary Table 18; Supplementary Data 10**). Spermidine level was down-abundant in BMA64-1A, BY4742, BY4741, and up-abundant in X2180-1A, SEY6210, and S288C under EtOH stress (**Supplementary Data 8**). To assess whether these spermidine levels impact the growth, we cultivated the strains under their highest EtOH stress level with and without spermidine supplementation. The supplementation increased the maximum population for BY4742 and BY4741, whereas it decreased for X2180-1A, SEY6210, and S288C (**Supplementary Figure 6D-E**). Therefore, strains with a low spermidine abundance under stress improved the growth when supplemented and vice-versa (BMA64-1A is the only exception). Overall, all LTs have a better growth under a higher abundance of spermidine, while this effect is observed only for two HTs (**Figure 2D**). We highlight that the gene SGF29 (YCL010C), which may be related to the spermidine on growth (further discussed), is down-regulated only in BMA64-1A and not DE in other strains (**Supplementary Data 6**).

### 2. Cell biology assays

The EtOH stress significantly impacts cell viability (**Table 1**), in which the number of viable cells (LL) is reduced for all strains, albeit S288C presented the highest cell viability (**Figure 3A-C; Supplementary Data 14**). The percentage of viable cells in LTs compared to HTs (LT divided by HT) in the controls are similar (0.82), while this rate under treatment is higher (1.45), emphasizing the lower cell viability in HTs under stress (**Figure 3B**).

**Figure 3:**
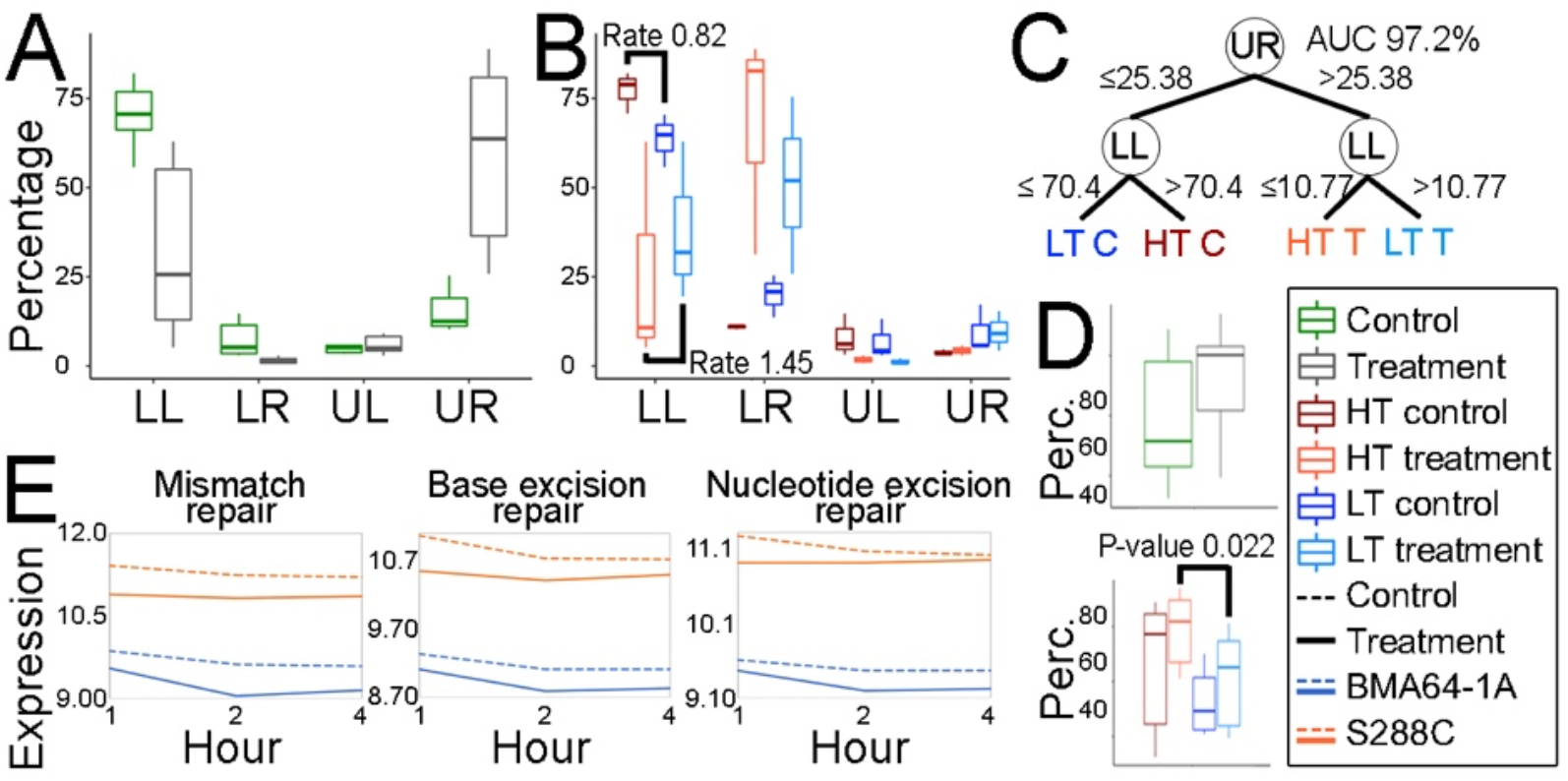
Cell viability, ROS accumulation, and analysis of DNA repair machinery. **A-B:** the percentages of viable cells (LL), cells under initial apoptosis (LR), and cell death (UL and UR). The “Rate” is the average of LT cells divided by the average of HTs. Hence, rate >1 indicates more LTs, whereas <1 is the other way around; **C:** decision tree to assess the relationship between LL and UR. The edges’ numbers are the percentage of cells under the condition branched to the adjacent node; **D:** ROS analysis; **E:** time-course expression landscape of genes related to the DNA repair.

The EtOH stress induced a higher accumulation of cells with ROS in HTs than LTs (rate LT/HT = 0.76, p-value = 0.02246), being BMA64-1A the most affected strain (**Table 3; Figure 3D; Supplementary Data 14**). Additionally, many genes related to oxidative stress were up-regulated in HTs (**Supplementary Table 15**).

**Table 3:** Fold-change of ROS, SDH, and DNA damage assays. **AVG:** average of replicates; **C:** control; **T:** treatment; *: statistically significant differences (p-value <0.05). Rate T/C >1 indicates the treatment’s value is higher than the control. Rate LT/HT >1 indicates that LTs have a higher percentage of cells under that condition.

| Group | ROS |  |  | SDH |  |  | DNA damage |  |  |
| --- | --- | --- | --- | --- | --- | --- | --- | --- | --- |
|  | AVG C | AVG T | Rate T/C | AVG C | AVG T | Rate T/C | AVG C | AVG T | Rate T/C |
| HT | 64.45 | 79.43 | 1.23 | 3.65 | 4.75 | 1.3 | 10.02 | 19.55 | 1.95 |
| LT | 52.52 | 60.39 | 1.15 | 2.88 | 3.25 | 1.13 | 8.46 | 21.57 | 2.55 |
| Rate LT/HT | 0.81 | 0.76* | - | 0.79 | 0.68 | - | 0.84 | 1.10 | - |

Mitochondria is the main source of intracellular ROS (Maris et al. 2001). Although there was a higher SDH activity in both phenotypes under EtOH stress, it was higher in HTs (see rate T/C) (**Table 1, 3**), fitting the ROS data. The up-regulation of mitochondrial division marker genes (e.g., FIS1, DNM1, and MDM36) for most strains (**Supplementary Data 6**) evidenced that the stress also induced mitochondrial division.

ROS can be a source of DNA damage (Mani and Chinnaiyan 2010). The EtOH stress induced a higher accumulation of cells with DNA damage in LTs than HTs (**Table 3; Supplementary Data 14**). Mismatch, base, and nucleotide excision repair genes (KEGG sce03440, sce03410, and sce03420) seem to be deeper down-regulated in S288C under treatment than control comparing to BMA64-1A, supporting the results of DNA damage assay (**Figure 3E**).

Since the flow cytometry data suggest that the DNA damage seems not to be repaired, we addressed this issue by assessing the expression of DNA damage checkpoint genes. The Rad53p and Chk1p branches are two mechanisms that simultaneously maintain the cell cycle arrest until the DNA repair: 1-Rad53p activates genes outcoming a cell cycle arrest; 2-activation of Chk1p causes phosphorylation of Pds1p also leading to cell cycle arrest (Sanchez 1999). RAD53 (YPL153C) is down-regulated in most strains, but only HTs presented a PDS1 (YDR113C) repression. Overall, only X2180-1A and BY4742 have both mechanisms hampered, indicating a normal cell cycle (**Supplementary Figure 34; Supplementary Data 6**).

Although nuclear RNA export, RNA synthesis, and degradation pathway genes were negatively affected by the EtOH stress, only in BMA64-1A a significant reduction of RNA yield in the whole-cell under stress was observed (**Supplementary Figure 22, 35**).

### 3. OMICs and network analysis

#### 3.1. Assemblings and differential expression

BMA64-1A genome sequencing had ~6,4M filtered reads with ~250 nts length, making up 134.88 X coverage on the final assembled genome. Only 36 coding genes did not have similarity to the S288C (**Supplementary Table 6**).

A total of 87-259 lncRNAs (~161 per strain) were found (Marques et al. 2021), ranging from ~200-400 nts. Most of these *loci* are near or within coding and intergenic regions in either sense or antisense orientation, there was no ‘intronic’ biotype (**Supplementary Table 8**), as expected (Yamashita et al. 2016; Novačić et al. 2020), and ~25.02% were classified as ‘others’. Small ORFs were found for ncRNAs in yeast (Smith et al. 2014). Although we found lncRNAs with small ORFs starting with methionine, the sequences did not match peptides from our proteomics and have no similarity to protein sequences neither domains. We observed few orthologues with ≥80% of similarity (**Supplementary Figure 7**). Additionally, the lncRNAs identified here have no similarity to those previously cataloged (Parker et al. 2017, 2018). Altogether, here is shown a *de novo* set of lncRNAs for yeast: GFF genomic coordinates are in https://figshare.com/account/home#/projects/115875.

Gene expression analyses were performed using our updated GFFs (https://figshare.com/account/home#/projects/115875). Around 50.31% and 47.76% of DEGs were up-regulated in HTs and LTs, respectively. The gene expression variation differs when comparing HTs vs. LTs (p-value = 0.0442), indicating the earlier have more up-regulated genes (**Supplementary Table 10**). A total of 1,330 and 868 significant DEGs (not considering lncRNAs) were observed within HTs and within LTs, respectively (**Figure 4A**). LncRNAs here found are less expressed than coding genes, as expected (Derrien et al. 2012; Djebali et al. 2012), albeit they tend to be up-regulated under stress (**Supplementary Figure 10; Supplementary Table 11**). Most TEs are up-regulated in both phenotypes under stress. Although LTs presented significantly higher TEs expression than HTs, the latter had a higher percentage of up-regulated ones (87.66% *vs*. 61.51%) (**Supplementary Data 7; Supplementary Figure 11**). All strains present a recent insertion wave of each repeat family and another smaller older wave older (**Supplementary Figure 8**).

**Figure 4:**
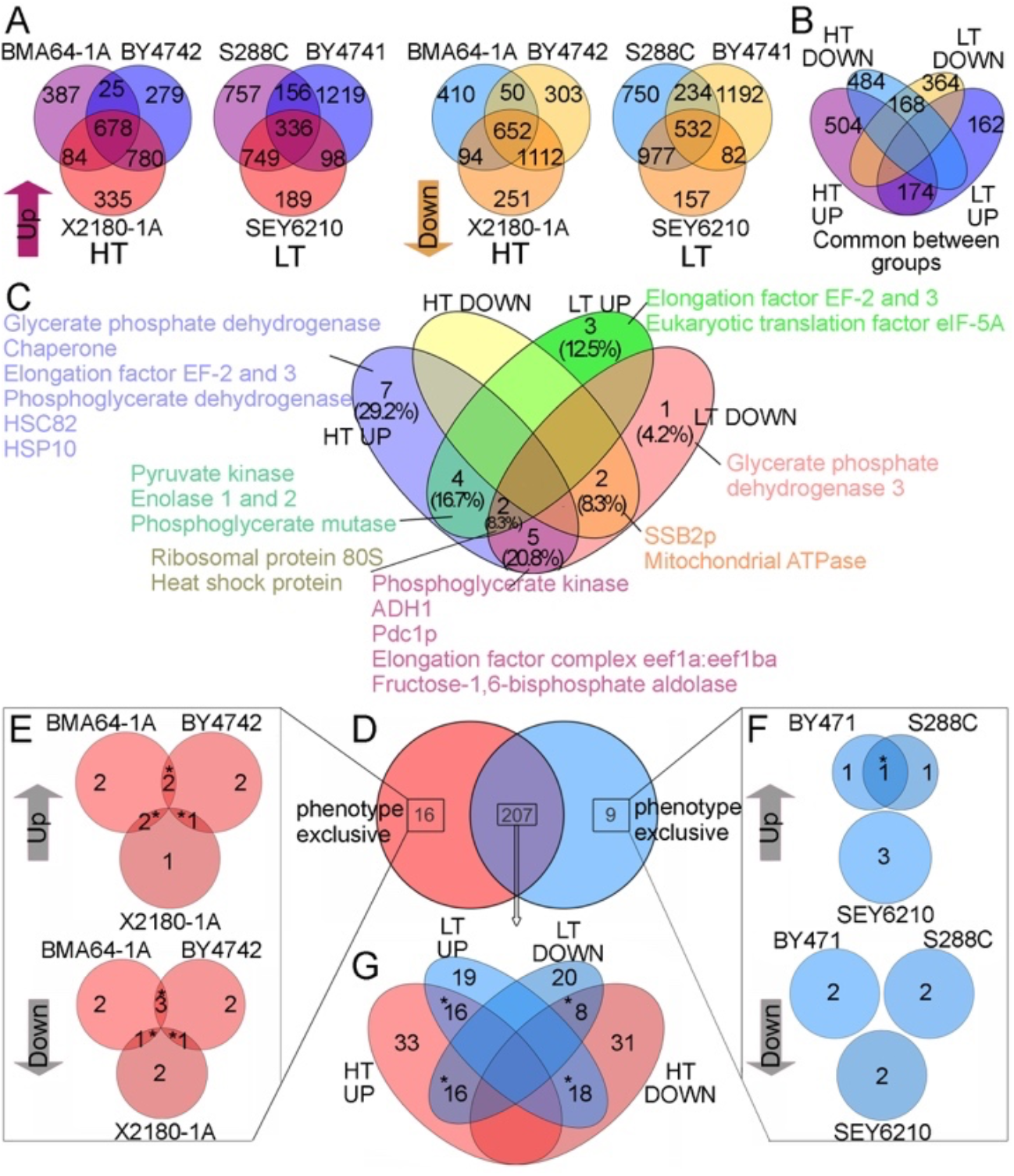
Differential expression analysis sets. **A-B:** DEGs amongst strains and phenotypes. In “B” is depicted sets between the up and down DEGs from the core of Venn diagrams in “A”; **C:** DAPs between phenotypes; **D-G:** DAMs amongst strains (“E”, and “F”) and phenotypes (“D”, and “G”). The “*” indicate DAMs mapped onto KEGG pathways; “Up” and “Down” are up-regulated and down-regulated, respectively.

Although many DEGs are shared between phenotypes (**Figure 4A**), they are not clearly expression-clustered (**Supplementary Figure 12**). However, functions shared between phenotypes were found. The up-regulated DEGs are related to metabolism, whereas most down-regulated are related to mRNA processing and ribonucleoprotein complex biogenesis (**Supplementary Table 17**). The GAGE-enrichment analysis of KEGG pathways showed that most DEGs in almost all strains are related to essential mechanisms, *e.g*., “citrate cell cycle (TCA cycle)” and “oxidative phosphorylation” (**Supplementary Table 18**). Phenotype specificities were found comparing the function between HTs’ and LTs’ genes (**Figure 4B**). The up-regulated genes in HTs are related to metabolism, while the down-regulated genes are related to essential processes such as DNA repair, division, and transcription. Either up or down-regulated genes in LTs are related to essential processes (**Supplementary Table 17-18**). We also found strain-specificities by another strategy (**Supplementary Table 15**).

A total of 20 and 19 DAPs were identified in HTs and LTs, respectively. We highlight the DAPs HSPs, ADH1, PDC1, elongation factors, ribosomal proteins, and chaperones (**Figure 4C**).

A total of 232 non-redundant DAMs were identified considering all samples. Most DAMs up-abundant in both HTs and LTs are related to TCA metabolism (**Supplementary Data 8, 10**), a pathway significantly disturbed by the EtOH stress according to our transcriptome GO enrichment analysis (**Supplementary Table 17**). Most DAMs (207) are shared between phenotypes (**Figure 4D**).

#### 3.2. Topological shifts of integrated networks

The 16 networks generated (6 strains + 2 phenotypes, for both control and treatment conditions) had a power-law degree distribution fitting the scale-free and Barabási-Albert models (Almaas and Barabási 2006). The networks were dissortatives, indicating resilience against random perturbations. The average path length (*d*) suggests a small-world effect. The EtOH stress increased the diameter (*dm*), betweenness centrality (*BET*), and Burt’s constraint (*BC*), and decreased the density, clustering coefficient (transitivity), the number of connections (*K*, or degree), and eigenvector centrality (*EIG*). The reduction of degree and the premature termination in the degree distribution tail under stress evidenced a loss of hubs (increased the intermediary ones): this is more intense in LTs. The normalized DDC function showed that the stress increased the probability of connection among nodes with a lower degree (**Supplementary Figure 16-17, Supplementary Table 13-14**).

#### 3.3. Defining function and structural analysis of EtOH responsive lncRNAs

The inferred LNCPI integrated most of the lncRNAs, averaging ~26.5 lncRNAs, ~207.16 target proteins and ~331.83 edges per strain, and ~8.1 target proteins per lncRNA (Marques et al. 2021) (**Supplementary Table 16; Supplementary Data 11**). These nets are reliable, fitting the number of edges (ncRNA-protein interactions) for yeast (Panni et al. 2017) and the number of target proteins (~5-20 per small RNA) (Chujo et al. 2016).

LncRNA functions assigned using guilt-by-association approach showed that most of the lncRNAs are related to “RNA polymerase”, “negative regulation of protein kinase activity by protein phosphorylation” (which includes RNA polymerase assembling mechanisms), “alcohol catabolic process”, and “telomere establishment processes” (**Supplementary Figure 18A**). A total of 8 and 10 target proteins were present only in HTs, and LTs, respectively. The roles of 4 targets shared amongst all strains evidenced a diversity of lncRNA’s functions, and the ones are not strictly related to the EtOH response (**Supplementary Figure 18B-E**).

LncRNAs targeting the same protein within S288C and within BMA64-1A (the intrastrain comparison) presented low similarities (~50.87%, and ~63.76%), and generally a lack of secondary structure conservation, such as expected for this RNAs (Derrien et al. 2012; Djebali et al. 2012; Nitsche and Stadler 2017). However, secondary structure conservation amongst orthologues to three S288C’s lncRNAs (≥80% similarity) was found in the interstrain comparison (**Supplementary Figure 7B-C, 19**).

The lncRNA-propagation analysis from each DE lncRNA showed that the up and down-regulated ones work on different systems within each strain. However, it was possible to cluster them into four functional categories: 1-“essential processes”, including transcription, replication, and ribosomal biogenesis; 2-“membrane-dependent processes”, including signaling and cell division, membrane/cell wall, and intracellular transport; 3-“metabolic processes”, including oxidative stress response, diauxic shift, fermentation, and other metabolisms; 4-“degradation process”. We highlight the interactions transcr_6448-GLN3 and transcr_18666-ATH1 (**Figure 5**). The time-course expression showed that under stress, transcr_6448 in BMA64-1A seems to cease the GLN3 (YER040W) increasing (negative correlation with R^2^=0.98), which is released when this lncRNA is significantly reduced; GLN3 is down-regulated only in BMA64-1A. Similarly, the negative correlation (R^2^=0.99) between the ATH1 (YPR026W) and transcr_18666 in S288C shows that ATH1 expression is increased when IncRNA is reduced; ATH1 is up-regulated in almost all strains (**Supplementary Figure 20B-C; Supplementary Data 6**).

**Figure 5:**
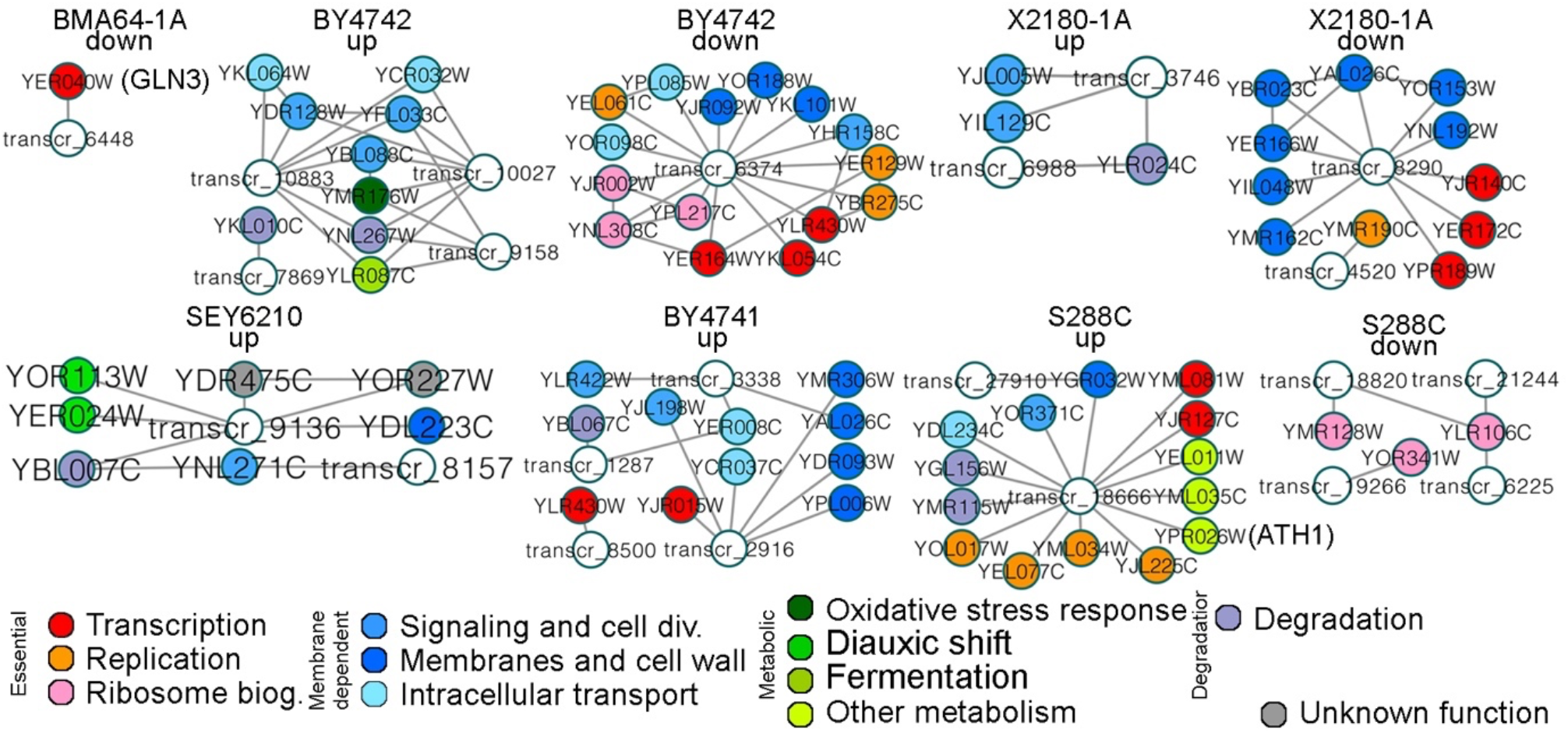
Analysis of LNCPI subsystems selected by lncRNA-propagation analysis. The node colors are related to the biological functions depicted at the picture bottom.

#### 3.4. Narrowing down the pathways affected by the EtOH stress

The transcriptome (1,175 DEGs), metabolome, and LNCPI networks were integrated onto 95 KEGG-maps per strain (https://figshare.com/account/home#/projects/115875). Almost all mapped lncRNA presented an inverse expression pattern to their target protein.

The most affected pathways were ranked based on DEGs, DAMs, DAPs, enrichment analysis, and lncRNA function assignments (**Supplementary Figures 13, 18, 21-22, 26, 28; Supplementary Data 6, 8, 10-12**). The EtOH stress-responsive molecules of pathways bellow mentioned will be appropriately called along the discussion to clarify our models. Details of these molecules are in **Supplementary Text** topics “4.3.2.”, and “4.3.3”.

There are 11 very affected life basal pathways: TCA cycle (TCA), glycolysis and gluconeogenesis (Gly/gluc), autophagy, MAPK, longevity, protein process in endoplasmic reticulum (PPER), RNA transport, ribosome biogenesis, mRNA surveillance, RNA degradation, and cell cycle, hereafter referred to node-pathways. Most DEGs of these pathways were down-regulated, although longevity, PPER, autophagy, TCA, Gly/gluc are exceptions: most of these induced genes are related to chaperones, heat shock proteins (HSPs are also up-abundant in our proteomics), and RNA and protein degradation systems (**Supplementary Figure 22; Supplementary Data 12; Figure 4C**).

The pathway integrated network (PINET) gathered the node-pathways in four communities: 1-MAPK and cell cycle; 2-TCA and Gly/gluc; 3-longevity, autophagy, and PPER; 4-mRNA surveillance, RNA degradation, RNA biogenesis and transport, and ribosome biogenesis (**Supplementary Figure 23A**). The EtOH stress caused network rewiring comparing strains, reported by our clustering analyses based on time-course data of BMA64-1A and S288C under stress (**Supplementary Figure 23B**; https://figshare.com/account/home#/projects/115875). Overall, the EtOH affected the MAPK, RNA-related pathways, ribosome biogenesis, and cell cycle (**Supplementary Figure 23B-24**).

The time-course expression landscape of cells under stress mapped on networks showed that peroxisome, Gly/gluc, and longevity pathways had an “up and stable” and “stable” profile in BMA64-1A and S288C, respectively. The landscape of RNA biology pathways is “down and stable” for BMA64-1A and “stable” for S288C. The TCA, cell cycle, MAPK, autophagy, and PPER presented concordant profiles between strains: the TCA (“up and stable”) is the only one from this set without a steady-state profile (**Figure 6A-B; Supplementary Figure 24**).

**Figure 6:**
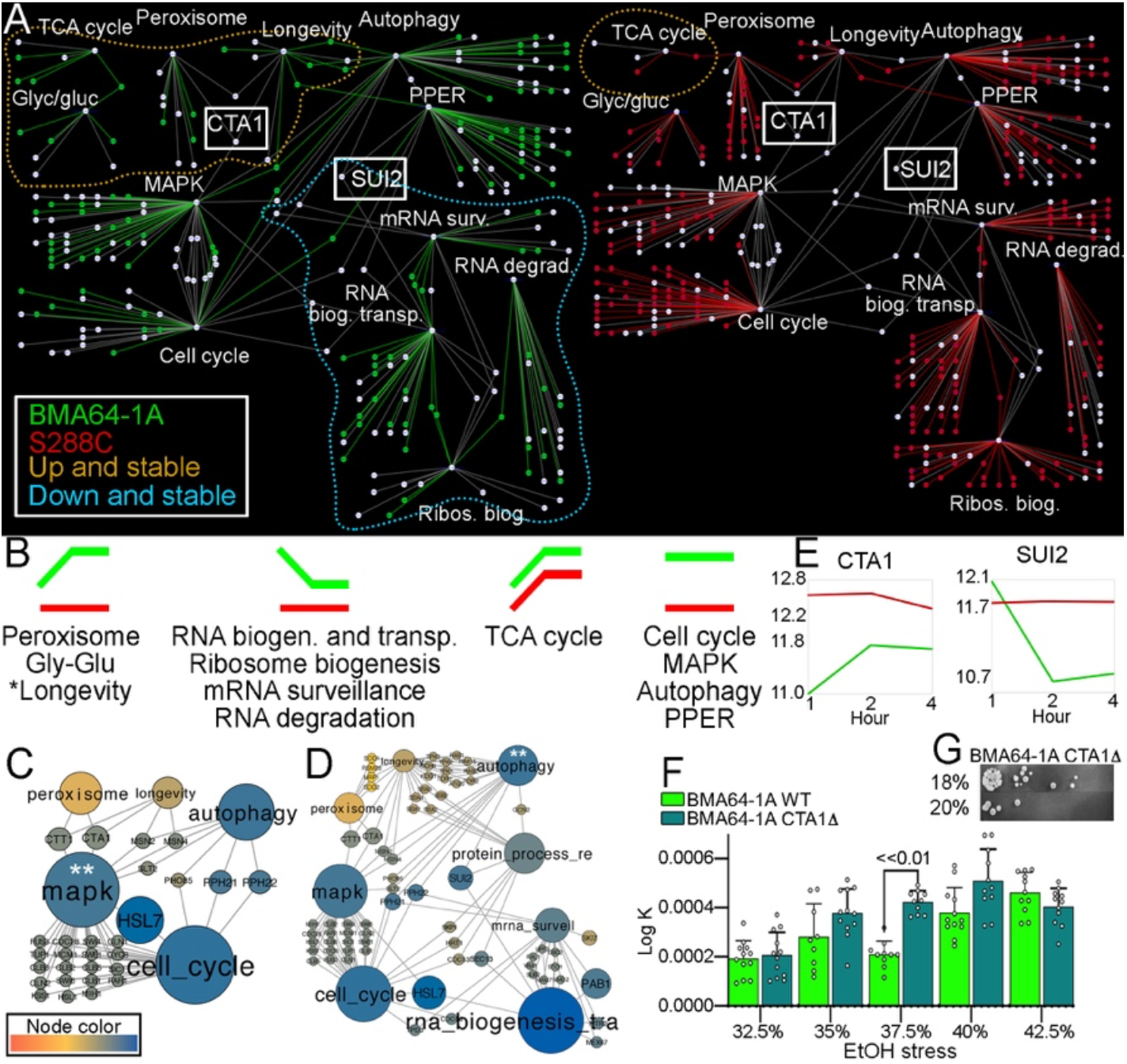
Life basal subsystems evidencing phenotype-divergences using BMA64-1A and S288C as models. Green and red colors depict the BMA64-1A and S288C data, respectively. **A:** comparison between networks modeled based on the time-course clusters (**Supplementary Figure 23B**). The white nodes and edges are the ones shared between BMA64-1A and S288C nets. Dot line delimits communities with similar time-course landscape; **B:** the overview of time-course landscapes from data in “A” under stress; **C-D:** network propagation analysis from node-pathways labeled by “**”; **E:** time-course landscapes of the crucial genes we claim to trigger the expression phenotype divergences under stress; **F:** population rebound after stress relief of BMA64-1A wild-type and CTA1Δ; **G:** spot test of BMA64-1A CTA1Δ. Details of “A”, and “C-D” are in **Supplementary Information 2** (https://figshare.com/account/home#/projects/115875).

To assess which pathway drives the mentioned divergences between phenotypes, we compared the expression landscapes (**Figure 6B**) *vs*. the network propagation (**Supplementary Figure 25**). We first inspected the subsystems from the MAPK pathway because it regulates the metabolism bridging external and internal signals (Chen and Thorner 2007). The data showed a direct information flow from MAPK to longevity, peroxisome, autophagy, and cell cycle (see the node-genes CTT1, CTA1, MSN2, MSN4, SLT2, and HSL7 in **Figure 6C**). Interestingly, the longevity and peroxisome are the first paths to present significant expression divergences (**Figure 6C**). Additionally, the upregulation and lack of DE for the longevity-related genes GPR1 (YDL035C), RAS2 (YNL098C), CYR1 (YJL005W), TPK1 (YJL164C), RIM15 (YFL033C), GIS1 (YDR096W), SOD1/2 (YJR104C/YHR008C), and CTT1 (YGR088W), most of the peroxisomal genes, and autophagy-related genes, evidenced the importance of these pathways. Moreover, the peroxisomal gene PEX12 (YMR026C) is only down-regulated in LTs, while the longevity-related gene SCH9 (YHR205W) is prevalently down-regulated in HTs (**Supplementary Data 6, 12**). The information further diffuses from autophagy mainly to RNA biogenesis (see the node-gene SUI2); interestingly, RNA biology genes in BMA64-1A presented an intense expression declining from 1h to 2h (**Figure 6B, D; Supplementary Figure 24C**). Finally, the information flows from longevity to peroxisome, autophagy, and PPER (many genes provide these links), as well as from peroxisome to TCA (**Supplementary Figure 25**).

The CTA1 (YDR256C) time-course landscape under treatment fits the ones of MAPK downstream-pathways longevity and peroxisome (**Figure 6A-C, E**). Then, we analyzed the population rebound and EtOH tolerance of BMA64-1A CTA1Δ to assess whether this gene is a regulator of phenotypes. Indeed, the highest EtOH tolerance level decreased in BMA64-1A CTA1Δ (20% v/v). This mutant had a better population rebound after stress relief than wild-type in most EtOH percentages tested, albeit only the difference at 37.5% is statistically significant (**Figure 6F-G**). These data corroborate our model: the EtOH tolerance is negatively related to the population rebound.

The SUI2 (YJR007W) time-course landscape under treatment fits the one of autophagy downstream-pathway RNA biology. The RNA biogenesis and transport pathway bridges other RNA biology pathways, which diverge between BMA64-1A and S288C (**Figure 6A-B, D**). Corroborating the BMA64-1A profile (“down and stable”), the acridine orange assay showed that only this strain has a significant RNA yield decreasing in the whole-cell under treatment (**Supplementary Figure 35**).

#### 3.5. Diauxic shift and the ethanol buffering

We modeled an EtOH stress-buffering mechanism based on the diauxic shift. Most genes in this model were up-regulated and impacted the metabolite abundances (**Figure 7; Supplementary Data 10**). Furthermore, the analysis of external glycerol yield (**Figure 2G**) also supports this model (further discussed). We highlight the up-regulation of genes essential for growth on non-fermentable carbon sources and diauxic shift responsive genes (NQM1 (YGR043C), CAT8 (YMR280C), ADR1 (YDR216W), INO4 (YOL108C), GUT1/2 (YHL032C/YIL155C), and RSF1 (YMR030W)) (Dasgupta et al. 2002; Klein et al. 2017), genes essential to convert acetyl-CoA in cytosol and peroxisomes (ALD4 (YOR374W), FAA1 (YOR317W), FAA2 (YER015W), HFD1 (YMR110C), ACS1 (YAL054C), PXA1 (YPL147W), PXA2 (YKL188C), POX1 (YGL205W), FOX2 (YKR009C), and POT1 (YIL160C)), carnitine acetyltransferases (CAT2 (YML042W), YAT1 (YAR035W), YAT2 (YER024W)), and TCA genes (CIT2 (YCR005C), GDH3 (YAL062W), GDH2 (YDL215C), KGD1 (YIL125W), and KGD2 (YDR148C)). Finally, we also found lncRNAs working on these pathways (**Figure 7; Supplementary Data 6**).

**Figure 7:**
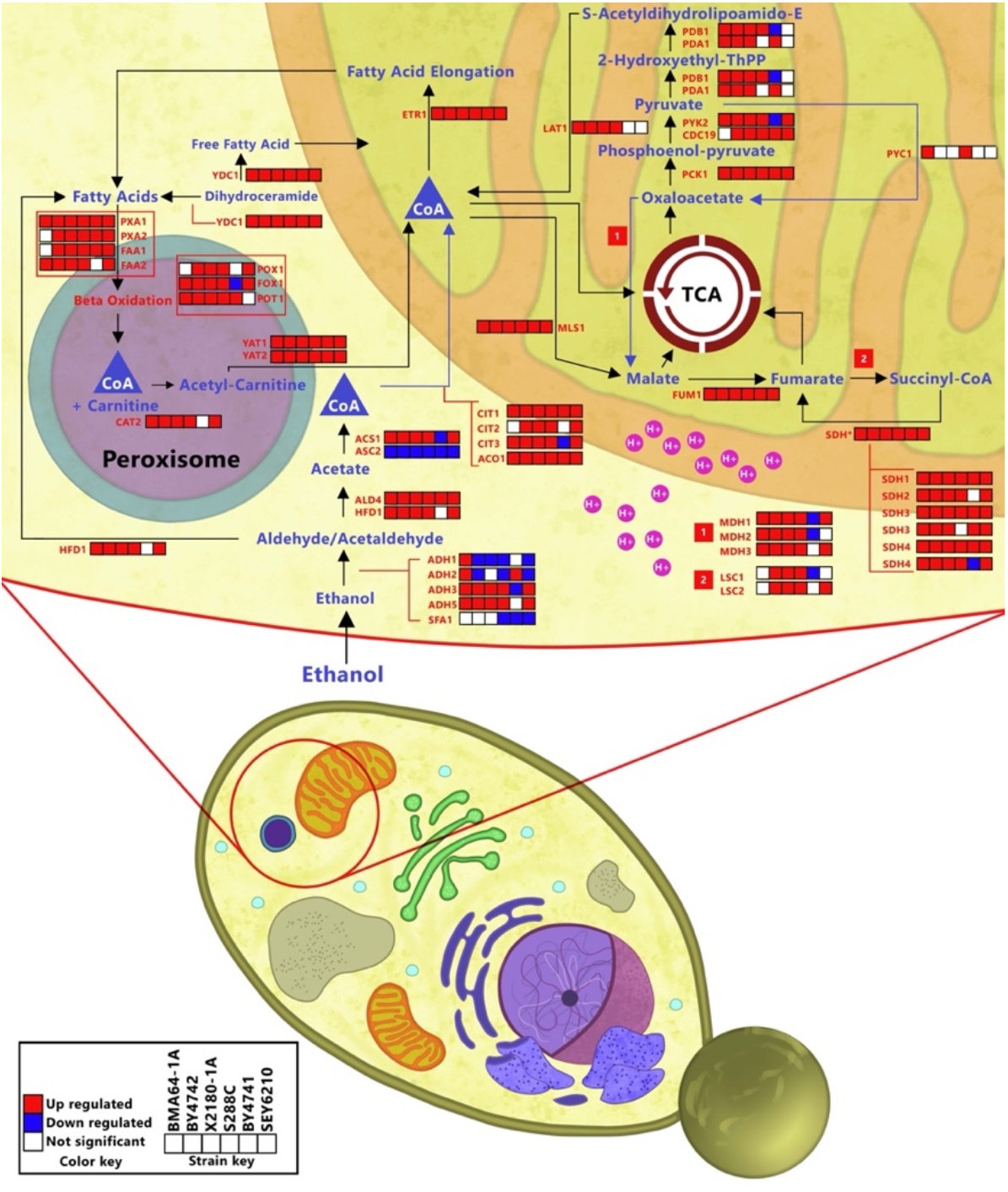
EtOH stress-buffering model by diauxic shift. Supplementary Information 3 (https://figshare.com/account/home#/projects/115875).

Transcr_20548 of BMA64-1A binds to proteins related to “positive regulation of transcription from RNA polymerase II promoter in response to EtOH”. The subnetwork harboring this lncRNA brings diauxic shift related genes (**Figure 8**). The time-course profiles and network edges diverged between BMA64-1A and S288C subnetworks, mainly concerning the ADH2. For instance, some BMA64-1A genes share similar transcriptional profiles, albeit those presented different profiles in S288C (**Figure 8A**).

**Figure 8:**
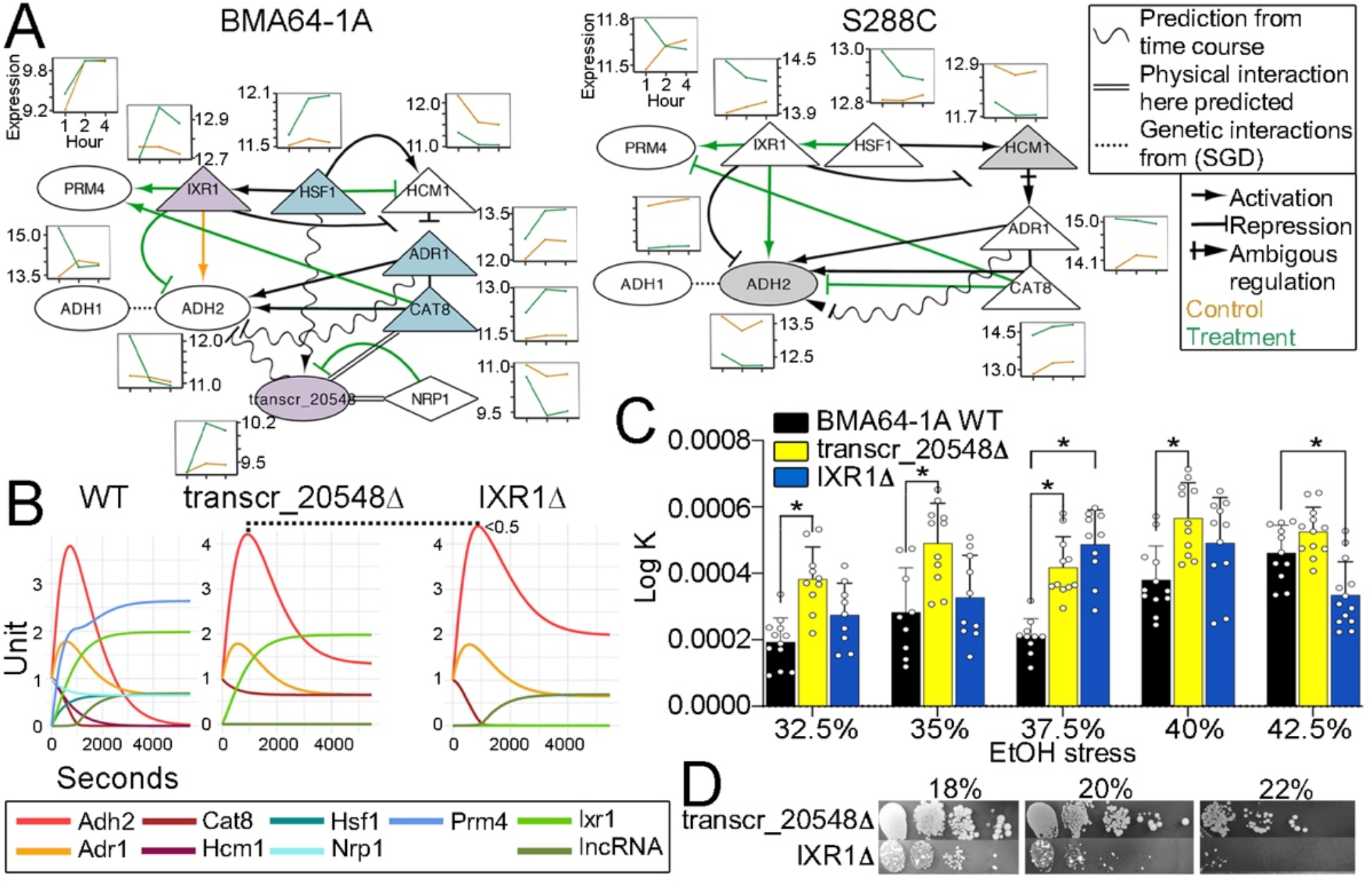
Subsystems model harboring the lncRNA transcr_20548, the diauxic shift, and EtOH stress-buffering genes. **A:** BMA64-1A and S288C subnetworks. Small boxes close to genes are the time-course data. Nodes sharing the same color have a similar time-course profile; **B:** BMA64-1A subnetwork dynamic model by ODE using the time-course of treatment condition. It simulates the gene expression of virtuals BMA64-1A wild-type (WT) and knock-outs (BMA64-1A transcr_20548Δ, and BMA64-1A IXR1Δ); **C:** population rebound after stress relief in WT, and CRISPR-Cas9 mutants. “*” is p-value <0.01; **D:** the EtOH tolerance spot test.

The ADH2 expression sharply decreased in BMA64-1A under EtOH stress. The ADH2 profile is negatively related to the ones in IXR1, ADR1, CAT8, and transcr_20548. Moreover, expressions depicted that IXR1 (YKL032C) and HSF1 (YGL073W) repress both HCM1 (YCR065W) and ADH2 (YMR303C), peaking the ADR1 (YDR216W), CAT8 (YMR280C), and lncRNA transcr_20548 expression under stress. Precisely, IXR1 and transcr_20548 expression profiles in BMA64-1A indicate that both seem to repress ADH2 under stress, which is not observed for S288C. The NRP1 (YDL167C) physically interacts with transcr_20548, probably negatively affecting the latter under EtOH stress (**Figure 8A**). Concerning the transcr_20548 regulation, its upstream genomic region has 12 motifs for the HSF1 domain (**Supplementary Figure 28D**), and the latter seems to act as a positive regulator of this lncRNA in both conditions (**Figure 8A**).

To assess whether the transcr_20548 of BMA64-1A is a stronger ADH2 repressor than IXR1, we evaluated the subnetworks dynamic using ODE and the population rebound of mutants. ADH2 enhanced in both transcr_20548Δ and IXR1Δ virtual mutants. Although this enhancement is slightly higher in IXR1Δ virtual mutant (~0.5 units), the IXR1 level is intrinsically higher than transcr_20548 (>2 units) in the virtual wild-type. Although previous findings showed that ADR1 and CAT8 synergically work on ADH2 expression (Walther and Schüller 2001), this was not observed in our time-course nor from simulated ADR1 (its level did not change comparing simulations). The CAT8 level decreases to ~0 in virtual wild-type and IXR1Δ, albeit it presents a steady-state (~0.6) in the virtual transcr_20548Δ. Furthermore, the ADH2 profile did not reach a steady-state in any simulation. Altogether, the ADH2 expression profile in the virtual transcr_20548Δ is not related to ADR1 and CAT8 expressions, showing they did not influence the full ADH2 regulation (**Figure 8A-B**). The population rebound after extreme EtOH stress showed that usually both BMA64-1A transcr_20548Δ and BMA64-1A IXR1Δ CRISPR-Cas9 mutants grown faster than the wildtype, and the earlier mutant often outperformed the IXR1Δ (**Figure 8C**). Both mutants reduced the EtOH tolerance to 20-22% (**Figure 8D**). The population rebound and spot test corroborate our model: the EtOH tolerance is negatively related to the population rebound.

#### 3.6. Membraneless structures affected by the EtOH stress

The transcriptome revealed that autophagy, P-bodies (PBs), RNA catabolic process, stress granules (SGs), proteasome stress granules (PSGs), proteasome and regulatory parts, protein polyubiquitination and positive regulation of ubiquitination (PPPR), and protein deubiquitination and negative regulation of ubiquitination (PDNR) are EtOH stress-responsive systems. A total of 191 genes related to the mentioned systems were selected from SGD to evaluate their expressions (**Supplementary Data 15**). We highlight that: 1-only few PBs-related genes were down-regulated (from 12.5% to 50%); 2-<50% of RNA catabolism process genes were down-regulated; 3-HTs presented a significantly smaller number of down-regulated genes related to the RNA catabolic process (T-test p-value=0.0241); 4-the BMA64-1A presented the highest percentage of up-regulated plus not DE genes for all pathways; 5-the BMA64-1A has the smallest percentage of down-regulated PB-related genes, whereas this percentage is similar amongst other strains (**Figure 9A**); 6-the decapping related genes DCS1 (YLR270W), DCS2 (YOR173W), and DCP2 (YNL118C) are not down-regulated in almost all strains; 7-essential genes of PBs and SGs (*e.g*., CDC33/eIF4E (YOL139C), eIF4A/TIF1 (YKR059W), DCP1 (YOL149W), DCP2 (YNL118C), PAB1 (YER165W), PUB1/TIA1 (YNL016W), SUP45 (YBR143C), and HBS1 (YKR084C)), mRNA decay and surveillance (*e.g*. LSM1 (YJL124C), DCP1 (YOL149W), DCP2 (YNL118C), RRP40 (YOL142W), RRP43 (YCR035C), RRP45 (YDR280W), RRP46 (YGR095C), RRP41/SKI6 (YGR195W), EDC3 (YEL015W), NUC1 (YJL208C), CAF40 (YNL288W), and MPP6 (YNR024W)), ribosome biogenesis and transport (*e.g*., CKA2 (YOR061W), CKB2 (YOR039W), RRP7 (YCL031C), UTP4 (YDR324C), UTP8 (YGR128C), GAR1 (YHR089C), EMG1 (YLR186W), NOP4 (YPL043W), among others) (Eulalio et al. 2007; Standart and Weil 2018; Falcone and Mazzoni 2018; van Leeuwen and Rabouille 2019) are not down-regulated in BMA64-1A, unlike almost other strains; 8-SGs-disaggregase genes HSP104 (YLL026W) and SSE2 (YBR169C) are up-regulated in almost all strains (**Supplementary Data 6**).

**Figure 9:**
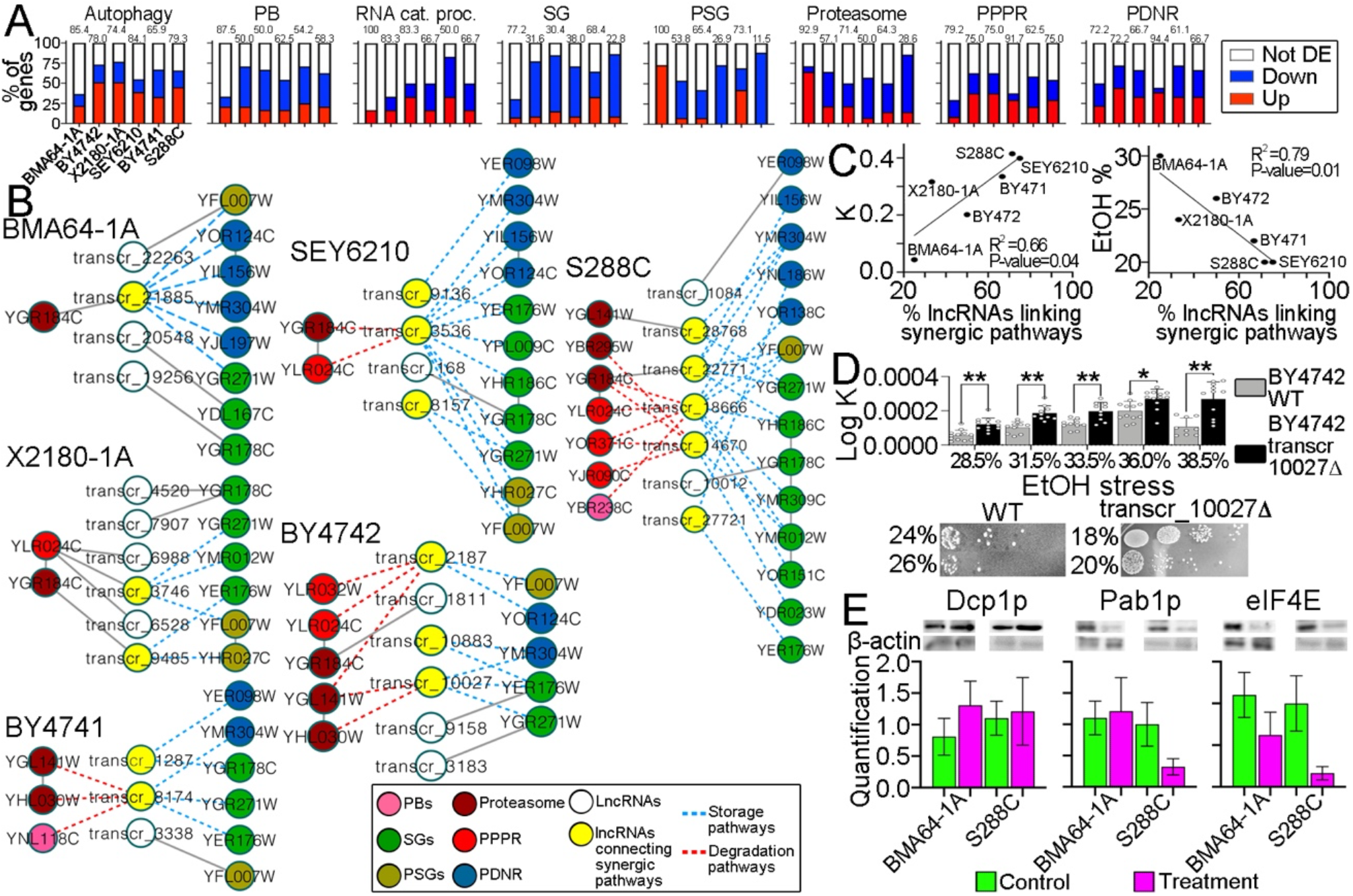
Analysis of genes and lncRNAs related to membraneless structures and degradation or storage EtOH stress-responsive systems. **A:** percentage of expressed genes. Numbers over the bars are the percentage of over-expressed plus not DEGs; **B:** subnetworks showing lncRNA-protein interactions related to degradation and storage systems; **C:** regression analysis comparing the percentage of lncRNAs connecting synergic pathways *vs*. the K parameter (growth rate) after the EtOH stress relief, or the percentage of the highest EtOH level *per* strain; **D:** the population rebound of BY4742 wild-type and the BY4742 transcr_10027Δ mutant, and their EtOH tolerance spot test. “*” and “**” are p-values <0.05 and <0.01, respectively; **E:** westernblot results.

The LNCPI showed that lncRNAs bind to 185 proteins related to “storage” (SGs, PSGs, and PDNR) and “degradation” (PBs, proteasome, and PPPR) pathways. The lncRNA-propagation analysis revealed the most relevant lncRNAs on these systems (**Figure 9B; Supplementary Data 15**). The percentage of lncRNAs connecting synergic pathways (within “storage” or “degradation”) is positively related to the population rebound after stress relief and negatively related to the highest EtOH levels analyzed (**Figure 9C**). Remarkably, the SEY6210 transcr_3536Δ was lethal, while the BY4742 transcr_10027Δ mutant had a reduced EtOH tolerance to 20% and an increase in the population rebound (**Figure 9D; Supplementary Figure 31F**).

The western blot showed an increase of Dcp1p and Pab1p and reduction of eIF4E in BMA64-1A under EtOH stress. Only Dcp1 p increased in S288C under stress. All these three proteins are higher abundant in BMA64-1A than S288C under stress, albeit the divergence is slight for Dcp1p and evident for Pab1p and eIF4E; only the Pabp1 difference is statistically significant (p-value=0.031) (**Figure 9E**).

## Discussion

According to the highest EtOH level to enable growth, we set the strains into HT (BMA64-1A, BY4742, and X2180-1A) or LT (SEY6210, BY4741, and S288C) phenotypes. Then, the highest EtOH level of a given strain is its most severe stress supported. We found that higher and lower EtOH tolerance phenotypes are neither related to the cell viability nor the faster population rebound after stress-relief; LTs presented the highest cell viability and population rebound. This rejects the hypothesis that a faster rebound counterbalances the high cell death rate from EtOH stress promoting a high EtOH tolerance phenotype; our mutants corroborated this conclusion. Therefore, LTs’ features here assessed may represent a higher adaptive response to the EtOH stress, as previously reported (Chi and Arneborg 2000).

We focus on identifying phenotype divergences, but strain-specificities were also covered. For some analyses, BMA64-1A and S288C were set up as HT and LT phenotype models, respectively. Different analyses showed a combination of pathways affected by the EtOH stress. Relevant pathways were selected based on the DEGs, DAMs and DAPs, GO analysis, KEGG mapping, and functional assignments of lncRNAs. The discussion bellow was split into five sections, and additional details are cited as **Supplementary Text** when needed.

The first section reports the EtOH stress-responsive lncRNAs. The key points are: 1-lncRNAs interconnect essential systems’ modules in a strain-specific manner to overcome EtOH stress; 2-although the EtOH stress-responsive lncRNAs are functionally strain-specific, they could be categorized into “essential”, “membrane-dependent”, “metabolic”, or “degradation” processes; 3-the NRP1 (YDL167C) repression reliefs the transcr_20548 of BMA64-1A from SGs under stress allowing this lncRNA to exert stronger repression than IXR1 on ADH2. This process outcome the low population rebound after EtOH stress relief observed for BMA64-1A; 4-the lncRNAs transcr_20548 and transcr_6448 of BMA64-1A are critical for its highest EtOH tolerance; 5-the highest population rebound and cell viability observed for S288C may be related to the lncRNA transcr_18666 action on the trehalose accumulation.

The second section reports the role of membraneless organelles, storage and degradation pathways in EtOH tolerance, and the involvement of lncRNAs on these systems. The key points are: 1-EtOH stress affords a suitable environment for membraneless structures; 2-cells under EtOH stress uses membraneless structures, and RNA/protein storage and degradation systems to endure the DNA damage harming; 3-HTs (mainly the BMA64-1A) take advantage of the degradation and storage processes; 4-the relationship between the number of lncRNAs connecting synergic systems and EtOH tolerance is still an unclear mechanism; 5-the transcr_3536 of SEY6210 and transcr_10027 of BY4742 are critical for their EtOH tolerance.

The third section concerns the information flow from EtOH stress throughout life basal systems. The key points are: 1-cells under EtOH stress keep active only basal pathways and tighten the systems to preserve and prepare themselves for the stress relief; 2-EtOH push intensive rewiring in life-essential subsystems, and the signals flow primarily from MAPK to longevity, peroxisome, and autophagy pathways; 3-the ROS accumulation and EtOH tolerance phenotype are early processes under EtOH stress; 4-CTA1, longevity, and peroxisome are the first EtOH-stress responsive mechanisms to exhibit phenotype-specific expression profile, being the longevity the primer one; 5-the repression of PEX12 (YMR026C) and SCH9 (YHR205W) seems to be related to the low EtOH tolerance in LTs and the higher expected oxidative stress resistance in HTs, respectively; 6-the response of autophagy-related genes under stress suggests a lifespan promote; 7-SUI2 (YJR007W) seems to bridge the network information flow from autophagy to RNA biology pathways, which is related to the RNA yield reduction observed in BMA64-1A under stress.

In the fourth section, we scrutinize our EtOH buffering model. The key point is that the diauxic shift drives an EtOH buffering mechanism, promoting an energy burst to hinder damages from this stress. HTs seem to take advantage of this process by reducing the oxaloacetate levels, inducing a higher SDH activity.

The fifth section reports the role of spermidine in the EtOH tolerance and the impact of EtOH stress on lipids metabolism. The key points are: 1-spermidine is critical for cell growth under the EtOH stress in a complex balance with the SGF29 (YCL010C) gene; 2-the sphinganine overload seems to be related to the lower cell viability under stress observed in HTs. It exacerbates the negative effects from the reduction of sphingolipids and inositol phosphorylceramide synthesis; 3-the inositol 1-phosphate accumulation in LTs may be an alternative to skip the harm from the low sphingolipid yield. This mechanism seems to be related to the higher cell surveillance, population rebound, and lower ROS accumulation observed for LTs; 4-the squalene and ERG9 (YHR190W) accumulation in HTs under stress seems to be related to their lower growth and cell viability under stress.

Finally, the putative role of eisosomes in cell acidification is speculated in “6. Other interesting genes and mechanisms” in the **Supplementary Text**. As far as we know, here is the first mention of this structure regarding the EtOH stress.

### 1. The EtOH stress-responsive lncRNAs are functionally diverse and likely involved in the EtOH tolerance

The number of lncRNAs and information regarding lncRNA-protein interactions on yeast is still scarce, and no reports associate them with EtOH stress. Herein, we present the largest catalog of *de novo* yeast’s lncRNAs for 6 strains focusing on the role of lncRNA-protein interactions on the EtOH stress. Overall, we found that the EtOH stress-responsive lncRNAs act as baits, backbones, or adapters interconnecting essential systems’ modules in a strain-specific manner to overcome stress challenges. Despite strain-specificities, we could define lncRNAs into broader categories mainly involving cell growth, oxidative stress response, degradation pathways, and diauxic shift. Additional details are in the discussion of **Supplementary Text**.

The lncRNAs found here presented canonical features such as an absence of *bona fide* ORFs, low sequence conservation, lower expression than coding genes, and the expected yeast lncRNA genomic organization and the number of ncRNA-protein interactions. The lncRNAs identified here are different from those already described (Parker et al. 2017, 2018). Finally, most of them are EtOH-stress responsive, corroborating that yeast lncRNAs are often responsive to environmental changes or stresses (Falcone and Mazzoni 2018).

LncRNA also works on proteins beyond gene regulation, *e.g*., acting on heterochromatin formation (Wang and Chang 2011; Niederer et al. 2017; Till et al. 2018). Additionally, stress response-related proteins usually are targeted by lncRNAs (Lakhotia 2012). Therefore, the guilt-by-association and the information flow throughout LNCPI (the lncRNA-propagation analysis) evidenced the lncRNA functions. Plenty of lncRNAs are related to the “RNA polymerase assembling”, “negative regulation of protein kinase activity by protein phosphorylation” (includes the “RNA polymerase assembling” plus “transcription by RNA polymerase II”), and “cellular alcohol catabolic process”. Considering only the EtOH stress-responsive lncRNAs, they act in a strain-specific manner by binding to proteins of a wide range of functions. However, we could classify these lncRNAs into categories “essential”, “membrane-dependent”, “metabolic”, and “degradation” pathways.s

Most lncRNA target-proteins related to “RNA polymerase assembling” are up-regulated in most strains, suggesting that the EtOH stress may induce the expression of RNAP-related genes. Interestingly, mutated Rbp7p (RNAPII subunit) enhanced the EtOH yield and tolerance, affecting the expression of glycolysis, fermentation, and oxidative stress response genes (Qiu and Jiang 2017); these pathways are prevalent in most of our analysis, corroborating these findings. Therefore, we modeled the role of lncRNA on RNAPII regulation using the lncRNA transcr_20548 of BMA64-1A as a model: the one works on the “positive regulation of transcription from RNA polymerase II promoter in response to EtOH”.

Our data suggest that under the EtOH stress and glucose starvation (**Figure 2G**), transcr_20548 may be contributing to the highest EtOH tolerance level and the lowest population rebound after stress relief observed for BMA64-1A. We suggest that Nrp1p seems to trap transcr_20548 into SGs, which is released after NRP1 repression. Hence, the strong repression of this lncRNA on ADH2 relies on NRP1 repression. Indeed, the BMA64-1A transcr_20548Δ overcame the IXR1Δ rebound after the stress relief and had a lower EtOH tolerance than wild-type; these data show the relevance of transcr_20548 on BMA64-1A EtOH tolerance. To understand the action of transcr_20548, we analyzed the timecourse and the dynamic system modeling. Firstly, we found that IXR1 negatively regulated ADH2, as previously reported for yeast under glucose starvation (Walther and Schüller 2001; Tachibana et al. 2005). However, transcr_20548 seems to be a stronger ADH2 repressor than IXR1. Unlike previous findings (Walther and Schüller 2001) neither ADR1 nor CAT8 influences ADH2 expression under stress at all. Secondly, we focus on understanding how the transcr_20548 regulation acts on ADH2. The transcr_20548 interacts with Nrp1p. The NRP1 time-course landscape is negatively related to the one of transcr_20548. Remarkably, Nrp1p (YDL167C) is a putative RNA-binding protein (Reynaud et al. 2001) and settles in SGs under glucose starvation (Buchan et al. 2008); SGs can link RNA-binding proteins (van Leeuwen and Rabouille 2019).

The time-course data and LNCPI showed that the lncRNA transcr_6448 of BMA64-1A might also help to induce its highest EtOH tolerance and lowest population rebound. This lncRNA seems to repress GLN3 (YER040W) under EtOH stress by some negative feedback loop. The GLN3 deletion boosts a branched-chain alcohol tolerance or may lead a cell cycle progression delay (Kuroda et al. 2019). We also hypothesize that GLN3’s regulator GCN4 (YEL009C) and YAP6 (YDR259C) may synergically work in this process since all these genes are down-regulated only in BMA64-1A.

The time-course data, LNCPI, and metabolome evidenced that the lncRNA transcr_18666 of S288C hinders Ath1p (YPR026W), resulting in a positive trehalose accumulation for S288C. The time-course data showed that ATH1 increases when transcr_18666 reduces. Ath1p degrades trehalose (Jules et al. 2004). Strikingly, the metabolome showed that trehalose increased in S288C even under ATH1 up-regulation, suggesting that transcr_18666 effectively hamper the Ath1p activity rather than its expression. Indeed, trehalose accumulation induces the EtOH tolerance and cell viability (Mansure et al. 1994; Lewis et al. 2010), as observed for S288C.

### 2. The membraneless organelles, storage, and degradation are related to EtOH stress: lncRNAs also act on these systems

The growth curves, flow cytometry, HPLC, transcriptome, LNCPI, and western blot, evidenced that the EtOH stress affords an environment to induce the storage and degradation pathways and the membraneless organelles. Some of these systems work on the degradation of defective RNAs and proteins produced from DNA damage. LncRNAs can scaffold RNAs and proteins creating membraneless structures (Pitchiaya et al. 2019; van Leeuwen and Rabouille 2019). Therefore, lncRNAs seem to have a joint action with the systems mentioned.

PBs degrade abnormal mRNAs and store RNAs to support long-term survival in stationary-phase (Eulalio et al. 2007; Ramachandran et al. 2011; Wang et al. 2018; van Leeuwen and Rabouille 2019). SGs improve the cell viability during stress by monitoring the stress, coordinating its relief, and precluding the mRNA degradation and translation initiation. SGs sequester mRNAs, translation initiation proteins, signaling proteins, and others, releasing them after stress (van Leeuwen and Rabouille 2019; Marcelo et al. 2021). PSGs protect cells from specific stresses, promoting a delay in cell re-entry into the proliferative state, genotoxic resistance, and fitness during aging. PSGs are proteasome reservoirs inhibiting its replenishment in the cytosol, which would degrade other proteins (Peters et al. 2013; Saunier et al. 2013). The autophagy system is essential to degrade damaged proteins and organelles, promoting cell survival under stress or starvation (Gatica et al. 2018). Overall, by either storing or degrading molecules, these systems are surveillance strategies until stress relief; hence, the shortage of these structures speeds up cell death under stress (van Leeuwen and Rabouille 2019). Besides the pathways mentioned, other RNA and protein degradation or storage systems are EtOH-stress responsive, as evidenced by the transcriptome and LNCPI. The ones include RNA catabolic process, autophagy, proteasome and regulatory parts, protein polyubiquitination and positive regulation of ubiquitination (PPPR), and protein deubiquitination and negative regulation of ubiquitination (PDNR). From now, we hereafter refer to the mentioned pathways and membraneless organelles (**Figure 9A**) as “storage and degradation systems”.

First, stationary-phase growth, oxidative stress, glucose starvation, and cytosol acidification under stress here observed are traits that afford a suitable environment for the assembling, or level increasing, of SGs, PSGs, and PBs (Eulalio et al. 2007; Buchan et al. 2008; Kato et al. 2011; Ramachandran et al. 2011; Shah et al. 2013; Wang et al. 2018; van Leeuwen and Rabouille 2019). Furthermore, as expected for cells accumulating SGs, and PSGs (Peters et al. 2013; van Leeuwen and Rabouille 2019; Marcelo et al. 2021), all strains here analyzed rebound after stress relief.

Second, we suggest that cells under EtOH stress seem to use membraneless structures and storage and degradation pathways to endure DNA damage harming, *e.g*., degrading defective RNAs and proteins resulting from that damage. Indeed, DNA damage enhances the RNA and protein mutants yield (Xie and Jarosz 2018). DNA repair mechanism abolishment relies on collapsing both RAD53 and CHK1-PDS1 mechanisms (Sanchez 1999). Although RAD53 and PDS1 expression suggest that 4 out 6 strains here analyzed had a cell cycle arrest by the DNA damage checkpoints, the flow cytometry showed DNA damage increasing in all strains under stress. Thus, we suggest that the presumed mutant RNAs and proteins resulting from DNA damage may be tackled by protective mechanisms induced by EtOH stress. Indeed, pre-treatment with EtOH enhanced the expression of genes related to protein quality control in yeast (Yoshida et al. 2020), and here the transcriptome suggests a normal function of RNA and protein degradation pathways: *e.g*., few genes related to PBs, RNA catabolic process, proteasome, and protein polyubiquitination were down-regulated. For instance, decapping genes DCS1, DCS2, and DCP2 are not down-regulated in most strains. Additionally, eIF5A, a PB’s protein (Moon and Parker 2018), is up-abundant in LTs’ proteomics, and many heat-shock proteins were either transcriptionally and translationally increased under stress. Since SGs prevent PBs to degrade mRNAs (van Leeuwen and Rabouille 2019), the observed higher down-regulation of SG-related genes than PBs suggests a certain PB activity level, hence, RNA decay.

Third, HTs (mainly the BMA64-1A) seem to take advantage of the degradation and storage systems. BMA64-1A presented the highest percentage of up-regulated (and lack of DE) for those systems-related genes; *e.g*., DCP1 is down-regulated in all strains except in BMA64-1A. Moreover, HTs had a significantly smaller down-regulation of RNA catabolism genes. Therefore, these data suggest regular or higher activation of these systems in HTs. To test this hypothesis, we compared the level of Dcp1p, eIF4E, and Pab1p between BMA64-1A and S288C. Dcp1 p and Pab1 p are present in PBs and SGs in yeast, respectively (Eulalio et al. 2007; van Leeuwen and Rabouille 2019). The 5’-3’ mRNA decay relies on the decapping mainly by Dcp1p and Dcp2p proteins (Parker 2012). The translation initiation factor eIF4E binds to the 5’ cap and joins with poly(A) tail binding protein Pab1p to enhance the translation initiation (Dever et al. 2016). However, the decapping and decay are independent of Pab1p presence since the ones also occur during translation reduction by decreasing the translation initiation factors (Parker 2012). The level of the mentioned proteins corroborates the decapping and translation stalling in both strains under stress, likely leading to mRNA decay. However, the higher protein levels in BMA64-1A under stress corroborates that this strain may take better advantage of these mechanisms, even though improving the protein holding into SGs.

Fourth, besides the lncRNA transcr_20548 of BMA64-1A (related to the SGs), we found four lncRNAs in BY4742 associated with the membraneless structures and degradation systems (details of transcr_10883, transcr_10027, transcr_9158, and transcr_7869 are in the **Supplementary Text** discussion). Therefore, we sought additional cases by reviewing our LNCPI and comparing this data with the gene expression of storage and degradation systems. By classifying lncRNA concerning linking synergic (within “storage” or “degradation”) or antagonistic (“storage” vs. “degradation”) systems, it shows that a smaller number of lncRNAs connecting synergic pathways seems advantageous to improve the EtOH tolerance, even though negatively impacting the population rebound after stress-relief. This hypothesis was tested by generating the BY4742 transcr_10027Δ and SEY6210 transcr_3536Δ mutants: the ones should have a lower population rebound and a higher EtOH tolerance. However, transcr_3536Δ is lethal, and the transcr_10027Δ mutant had a lower EtOH tolerance and a higher population rebound than wild-type. Although this data demonstrates that these lncRNAs are crucial for the EtOH tolerance of those strains, the relationship between lncRNA linking synergic pathways and EtOH tolerance is a complex mechanism not clarified here. On the other hand, the transcr_10027 of BY4742 binds to the Tel1p (YBL088C), and this gene represses the PBs formation (Tkach et al. 2012). Then, we speculate that the lack of this lncRNA might be releasing Tel1p blocking PBs. This process would speed up the transcr_10027Δ cell cycle re-entry after stress relief, such as observed.

Finally, we suggest that the cell cycle re-entry observed in the rebound experiments are SG-dependent mediated by the up-regulation of HSP104 (YLL026W) and SSE2 (YBR169C). These genes code proteins to dissolute the yeast’s SGs, allowing the reactivation of proteins and RNAs arrested inducing the cell cycle re-entry (Kroschwald et al. 2015).

### 3. EtOH causes intensive systems rewiring in life basal pathways: longevity pathway, CTA1, and SUI2 as master-key regulators

We show that EtOH stress prompts intensive systems rewiring keeping only basal modules active. The rewire of life basal systems longevity, peroxisome, Gly/gluc, and RNA biology pathways outcome expression phenotype divergences: longevity, peroxisome, CTA1, and SUI2 are master-key regulators.

Under the EtOH stress, cells maintain only the basal pathways active and tighten the whole system. We argue these features are cells mechanisms to preserve and prepare themselves for stress relief; our transcriptome also evidences the activation of basal modules. In this case, cells under EtOH stress redirect the systems’ signal by creating longer paths and signal delay to specialized late response pathways, driving activation of basal systems solely, evidenced by the increasing of diameter, path length, betweenness, and reduction of density and transitivity. Indeed, the low transitivity indicates that hubs’ neighborhoods are sparsely connected (Mason and Verwoerd 2007) suggesting longer paths. Longer paths favor the network modularity (Xu et al. 2011) leading to a signal delay from regulators to regulatory responses (Klein et al. 2012). We showed that EtOH stress reduced the alternative routes tightening the systems, evidenced by the decrease in the degree, eigenvector, and the connection of highly connected hubs (increases the connections of intermediary ones). These features indicate that less important nodes lose connection with more relevant ones (Lohmann et al. 2010). The increase in the Burt’s constraint and the decrease in density (such as here observed) evidenced a reduction in alternative routes availability (Burt 2004; Buskens and van de Rijt 2008). Therefore, we suggest that longer paths, systems’ delay, and systems-tightening prompt (or muses) lower flexibility on active pathways under EtOH stress. Altogether, cells would be preparing for a rapid and dynamic response from stimuli detected in the medium (Mangan et al. 2003).

To better understand the modification mentioned, we evaluated subsystems selected based on our transcriptome and proteome. Again, EtOH stress leads to intensive rewiring even in the basal-systems MAPK, longevity, autophagy, TCA, RNA-related pathways, RNA and protein degradation systems, Gly/gluc, ribosome biogenesis, cell cycle, and protein process in the endoplasmic reticulum (see **Supplementary Figure 23**). Overall, we claim that the EtOH tolerance phenotype and ROS accumulation are primary stress responses soon after signaling from MAPK to longevity, peroxisome, autophagy, and RNA biology pathways. Indeed, MAPK, ROS, and mitochondrial-related pathways have an important role against the EtOH stress (Li et al. 2019a).

Longevity, peroxisome, and CTA1 is the first responsive set to exhibit phenotype-specific gene expression, being longevity the trigger. Although MAPK regulates the metabolism bridging external and internal signals (Chen and Thorner 2007), this pathway did not exhibit expression divergences between phenotypes. The longevity path includes cell signaling proteins and, jointly with peroxisome, compose downstream elements from MAPK, mediated by the CTA1 gene, promoting substantial expression divergences between HTs and LTs. Precisely, the prevailing better population rebound in the CTA1Δ mutant here observed, corroborates the CTA1 as a master-key regulator of EtOH tolerance. Despite previous reports already showing the role of CTA1 in EtOH tolerance (Du and Takagi 2007) and chronological lifespan (Weinberger et al. 2010), here we pinpoint its primer role on outcoming these features by directly linking this MAPK stress signaling to longevity.

Besides the ROS measurement by flow cytometry, the expression of CTA1, SCH9, CTT1, and PEX12 are additional evidence that ROS accumulation is an early process of EtOH stress. The ROS accumulation increases the oxidative stress demanding higher oxidative stress resistance (Jamieson 1998). The lack of CTA1 reduces oxidative stress resistance (Okada et al. 2014). Thereby, the mentioned information flow mediated by CTA1 may also be a by-product of higher ROS accumulation observed for HTs, which would demand higher oxidative stress resistance: *e.g*., BMA64-1A had a CTA1 expression intensification under 2h of stress, besides the highest ROS rate. Moreover, the lack of SCH9 increases the cell resistance to oxidative stress (Teixeira et al. 2014). The SCH9 (YHR205W) expression dampening in all HTs may be related to their expected higher oxidative stress resistance. Such as here observed for all strains, oxidative stress and carbon starvation induce the expression of the longevity-related gene CTT1 (YGR088W) which deals with oxidative stress by inactivating ROS (Herrero et al. 2008; Auesukaree 2017). The peroxisome biogenesis (relevant source of ROS) requires the gene PEX12 (YMR026C) expression (Chang et al. 1997; Auesukaree 2017), and this gene is required for EtOH resistance (Teixeira et al. 2009). Therefore, the reduction of PEX12 observed for LTs may be responsible for their low EtOH tolerance and ROS. Finally, the up-regulation in bulky peroxisome-related genes evidenced the relevance of this organelle in the EtOH tolerance.

After information flow from MAPK to longevity and peroxisome, our data showed that signal flows toward autophagy and RNA biology pathways. Autophagy is related to cellular protection, promoting the lifespan through nutrient restrictions, oxidative stress response, DNA damage response, and metabolic changes (Longo and Fabrizio 2012). Therefore, the scarcity of down-regulation in the autophagy-related genes suggests that strains may be promoting a longer lifespan under the EtOH stress. Our data reported that SUI2 carries information from autophagy to RNA biogenesis and transport. Furthermore, SUI2 and RNA biogenesis and transport presented the time-course expression profile “down and stable” in BMA64-1A and “stable” in S288C, which may be related to the significant reduction of RNA yield BMA64-1A under stress. Although previous findings report that SUI2 increases the RNA accumulation in cells under high temperatures (Williams et al. 1989), as far as we know, here is the first evidence of a relationship between its expression and RNA accumulation under EtOH stress. Additionally, Sui2p is associated with the membraneless organelle eIF2B body (Moon and Parker 2018), and here we observed those structures are EtOH-responsive, as already discussed.

### 4. The ethanol buffering model

The transcriptome, metabolome, KEGG mapping, ROS measurements, glucose influx, and network analysis revealed that many pathways related to energy and detoxification were affected by the EtOH stress. Herein, there is an integrated view of both energy and detoxification systems in a model hereafter referred to as “EtOH buffering” (**Figure 7**). In this model, EtOH stress drives a diauxic shift by consuming EtOH and glycerol, intensifying the acetyl-CoA, fatty acids, TCA, and peroxisome activities outcoming an energy boost. Therefore, this mechanism would buffer the intense EtOH stress dampening its harms. Indeed, previous findings showed that a higher energy yield from the TCA helps to endure the EtOH stress (Lourenço et al. 2013). Finally, HTs seem to take better advantage of this process, fostering their higher EtOH tolerance.

*S. cerevisiae* under glucose depletion uses EtOH or glycerol as a carbon source (Klein et al. 2017). Our findings suggest that EtOH stress leads cells to the diauxic shift. We observed that EtOH stress prompts: 1-glucose starvation; 2-an up-regulation of diauxic shift responsive genes and essential ones for the cell growth on non-fermentable carbon sources (Dasgupta et al. 2002; Klein et al. 2017); 3-an apparent EtOH consumption. In this case, two alcohol dehydrogenases (ADHs which also catabolize EtOH, KEGG sce00010) are up-regulated. Quinolinate is increased under stress. This metabolite is essential for NAD+ synthesis, which in turn, is crucial for ADHs (di Luccio and Wilson 2008; Liu et al. 2019). Additionally, NADH helps in alcohol detoxification (Hasunuma et al. 2014); 4-a glycerol consumption to produce D-glyceraldehyde, evidenced by increasing of latter metabolite under stress and up-regulation of GCY1 (YOR120W catalyzes this reaction, (Klein et al. 2017)). However, only 3 strains presented glycerol retention even though the glycerol efflux controller FPS1 (YLL043W) is down-regulated in almost all strains; 5-an apparent lower acetaldehyde yield and EtOH buffering in LTs. EtOH negatively regulates PDC1 (YLR044C) (Liesen et al. 1996). Pdc1p converts pyruvate to acetaldehyde (KEGG sce00010) (Hohmann and Cederberg 1990), and the one is reduced and increased in LTs, and HTs, respectively. The alcohol dehydrogenase Sfa1p (YDL168W performs the reversible reaction from EtOH to acetaldehyde or aldehyde, EC number 1.1.1.1, KEGG pathway sce00010) is down-regulated only in LTs.

Acetaldehyde or aldehyde from EtOH catabolism would be converted to acetyl-CoA in cytosol and peroxisomes by different enzymes (Black 1951; Shani and Valle 1996; Hiltunen et al. 2003; Nakahara et al. 2012; Chen et al. 2012; de Jong et al. 2014) (details of reactions are in the discussion in the **Supplementary Text**). The genes involved in this pathway were up-regulated in most strains here analyzed.

Acetyl-CoA is transferred to mitochondria by four carnitine acetyltransferases (CATs). CAT Cat2p converts the acetyl-CoA in acetyl-Carnitine in the peroxisome (van Roermund et al. 1999), further transported to mitochondria by the CATs Yat1p and Yat2p. Interestingly, double deletion of CATs YAT2 and TCA cycle CIT2 do not allow growth on non-fermentable carbon sources mediums (Swiegers et al. 2001). According to our transcriptome analysis, HTs may be taking advantage of using acetyl-Carnitine since the carnitine metabolism is an enriched term for these strains. Interestingly, Yat2p binds to lncRNAs in BY4742, SEY6210, and BY4741, albeit only the interaction in SEY6210 is reliable according to the lncRNA-propagation (information flow analysis). However, the effects of lncRNA-YAT2 interactions are unclear. A surmise positive action on acetyl-CoA transport, lncRNAs would contribute to the energy boost during the severe EtOH stress. Acetyl-CoA in mitochondria also seems to trigger a positive feedback loop for its synthesis in the peroxisome through the fatty acid elongation pathway; this path is important for the EtOH tolerance (Lewis et al. 2010). The fatty acid elongation starts with the decomposition of acetyl-CoA mediated by Etr1p (YBR026C) (Miinalainen et al. 2003) (which is up-regulated in all strains here analyzed) (KEGG sce00062). This fatty acid backs to peroxisome and is converted to acetyl-CoA, as mentioned.

Acetyl-CoA enters into the TCA cycle in mitochondria to generate energy (Swiegers et al. 2001; Franken et al. 2008). Metabolomics, GWAS, and networks data evidenced the association between TCA and EtOH stress (Ming et al. 2019; Kang et al. 2019). Many data evidenced the activation of the TCA cycle in all strains analyzed here. For instance, this pathway is the only one to present a time-course profile “up and stable” in BMA64-1A and S288C. KEGG pathway enrichment showed the TCA cycle is induced in all strains under EtOH stress here analyzed. SDH assay also evidenced that acetyl-CoA enters in TCA cycle. The metabolome evidenced differential abundances in three relevant TCA metabolites (oxaloacetate, fumarate, and malate) under stress.

We suggest that HTs’ energy boost outcome from the lack of oxaloacetate and higher SDH activity along the TCA cycle. Acetyl-CoA condenses with oxaloacetate to form citrate (EC number 4.1.3.7). Conversely to LTs, the down-abundance of oxaloacetate in HTs indicates a lower acetyl-CoA catabolism in these strains. However, HTs take advantage of the lack of oxaloacetate. This metabolite is a potent SDH inhibitor (Priegnitz et al. 1973), and hence, its lack in HTs allowed for its higher SDH activity observed, probably outcoming an energy boost. In this case, higher SDHs activity increases the fumarate yield, rendering additional FADH2 for the mitochondrial respiratory chain to pump H^+^. The fumarate is further converted to malate (Martínez-Reyes and Chandel 2020). Remarkably, the expected higher abundance of fumarate in HTs is not observed, whereas its level in LTs did not change, as expected. However, the observed reduction of fumarate in HTs may be related to a putative high conversion of this metabolite to malate, which is indeed up-abundant in these strains. The strong up-regulation observed in bulky TCA enzyme coding genes in all strains is another evidence of either higher TCA cycle activity and H^+^ pumping mentioned. The role of these enzymes in NADH and H^+^ pumping is well known (Repetto and Tzagoloff 1989; Miller and Magasanik 1990; DeLuna et al. 2001; Reinders et al. 2007; Martínez-Reyes and Chandel 2020) (see **Supplementary Text** discussion).

Finally, both phenotypes seem to increase the mitochondrial division in an attempt to support EtOH stress, albeit HTs seem to take advantage of this process. The EtOH stress-induced the up-regulation of PHB1 in almost all strains and PHB2 only in HTs. These genes work on mitochondrial segregation and reduce senescence (Berger and Yaffe 1998; Piper et al. 2002).

### 5. The complex balance of spermidine and the impact of ethanol on lipid metabolism

Spermidine is crucial for cell growth under EtOH stress: the high abundance of spermidine allowed better growth for all LTs and two HTs. Previous findings showed that spermidine positively influences yeast growth in the presence of EtOH (Kim et al. 2017). Interestingly, the lack of spermidine decreases the lifespan and increases the ROS and necrotic levels (Eisenberg et al. 2009). Therefore, we suppose that spermidine may also be responsible for the highest cellular viability and lower ROS accumulation in LTs. Remarkably, the highest EtOH tolerant strain (BMA64-1A) is the only one in which spermidine level did not influence the growth. Further, we sought a gene that could be related to this BMA64-1A feature. The spermidine addition causes hypoacetylation in H3 lysine acetylation increasing the lifespan (Eisenberg et al. 2009), while SGF29 (YCL101C) acetylates the same residue (Bian et al. 2011): hence, the hypoacetylation and increasing the lifespan mentioned would rely on spermidine yield and a scarcity of SGF29. However, the data of BMA64-1A suggests a more complex scenario. In this case, the spermidine positive role on growth would depend on a certain level of H3 lysine acetylation since only this strain decreased both SGF29 and spermidine under stress: strikingly, this metabolite did not influence its growth.

Many papers showed the effects of EtOH stress on lipids. Remarkably, lipidomic analysis in yeast strains under different EtOH stress revealed differences in the lipids saturation rather than considerable differences in composition (Lairón-Peris et al. 2021). However, the transcriptome analysis here suggests that lipid metabolisms are altered in cells under EtOH stress. Such as previously found, many genes related to these metabolisms (*e.g*., ETR1, GPD1, MCR1, OPI3, FAA1, and GRE2) are induced under EtOH stress (Ogawa et al. 2000; Chandler et al. 2004). Then, here we addressed a deeper analysis of lipids metabolism comparing our transcriptome and metabolome seeking phenotype-specific divergences and specific patterns for BMA64-1A. From now, see the model in **Supplementary Figure 21B**.

Our data suggest that EtOH seems to reduce the sphingolipids (KEGG sce00600, mainly ceramides), and inositol phosphorylceramide (IPC) synthesis, which may be related to the decreasing of cell viability on both phenotypes under stress. Furthermore, we claim that the sphinganine enhancement observed only for HTs may be responsible for exacerbating the adverse effects mentioned; HTs present a lower cell viability and population rebound. In fact, the synthesis inhibition of sphingolipids, ceramides, IPC, and the lack of AUR1 harsh the cell division, growth, and viability (Wu et al. 1995; Giaever et al. 2002; Epstein et al. 2012; Katsuki et al. 2018).

In the sphingolipids/ceramides pathway, we observed the up-regulation of YDC1 (synthesize sphinganine). The over-expression of this gene reduces the growth, stress resistance and enhances apoptosis (Aerts et al. 2008; Yoshikawa et al. 2011), as observed here. Interestingly, the expected sphinganine up-abundance is observed only for HTs, which may be related to their lower population rebound after stress relief. Previous findings showed that sphinganine overloads inhibit the synthesis of sphingolipids and ceramides, hindering cell division (Wu et al. 1995; Epstein et al. 2012). On the other hand, there is a reduction of sphingosine in both phenotypes under stress and a high number of down-regulated sphingolipids/ceramides genes in all strains. PLC1 inactivation decreases the competitive fitness in a medium with EtOH (Qian et al. 2012). Plc1p synthesizes inositol 1-phosphate (myo-Inositol 1-phosphate or 1D-myo-inositol 1-phosphate), a substrate for Aur1p to produce IPC: it links IPC and sphingolipids/ceramides pathways (Becker and Lester 1980; Nagiec et al. 1997; Le Stunff 2002; Ramirez-Gaona et al. 2017). Despite PLC1 and inositol 1-phosphate levels increased only in LTs, the IPC synthesis may be compromised under EtOH stress since AUR1 is down-regulated in all strains. Altogether, sphingolipids, ceramide, and inositol phosphorylceramide (IPC) synthesis seem to be compromised under EtOH stress.

LTs seem to have an alternative pathway to skip damages imposed by the sphingolipids reduction by mean PLC1 and inositol 1-phosphate. Although inositol 1-phosphate enrichment in the medium improves neither EtOH nor cell viability (Ishmayana et al. 2020), the cell viability in LTs is higher than HTs. PLC1 inactivation delays the cell cycle progression (Lin et al. 2000), and its overexpression increases the level of other inositols (Tisi et al. 2004; York et al. 2005). Inositol-based metabolites are protective compounds under many abiotic stresses (Valluru and Van den Ende 2011). Altogether, the accumulation of PLC1 and inositol 1-phosphate in LTs may be improving their cell viability, population rebound, and lower ROS accumulation.

The accumulation of squalene (steroid metabolism, KEGG sce00100) and ERG9 (YHR190W) only in HTs under stress may be related to their lower growth and cell viability. ERG9 synthesizes squalene, and overexpressing this gene reduces the vegetative growth in yeast (Yoshikawa et al. 2011). Additionally, squalene accumulation negatively affects population growth and viability (Garaiová et al. 2014; Valachovic et al. 2016; Csáky et al. 2020).

## Acknowledgements

The authors thank Dr. Lucilene Delazari dos Santos for guiding us concerning the proteomics data, Mr. Edgar Allan Gobo for the artistic drawing of Figure 7, and Giltae Song for run the AGAPE pipeline to annotate the BMA64-1A genome.

## Author contributions

Main authors in the project conception: GTV, RMTG, LFA, IRW

Growth curves: LFA, LNM, LCL, AFJ, FBF, RMTG

OMICs data acquirement: LFA, CCOA, LHC, TC, LNM, MVL, CAL

Assembling and annotation: IRW, LFM, ER, RLBC, GTV

Differential expression: IRW, CCOA, LHC, RLBC

Cytometry, and microscopy: MAG, LAF, LNM, RTN, CCOA, LHC, LCL, APS, CM

Molecular biology: APS, LHC, MNF, LAJJ

Data integration and networks analysis: GTV, IRW, LFM, LCL

Influx and efflux analysis: CMP, JN, AD

Statistics: IRW, ROA, RPS, GTV

Main data interpretation: GTV, IRW, LFM, LFA, APS, LCL, LHC

Manuscript drafting: GTV, IRW, LFM, LCL

Final version: all authors

## Additional information

Supplementary data are available at BioProject number PRJNA727478, and https://figshare.com/account/home#/proiects/115875. Companion materials are Marques et al. 2021. “Long non-coding RNAs bind to proteins relevant to the ethanol tolerance in yeast: a systems biology view”. BioArxiv. https://doi.org/10.1101/2021.02.07.430053, and Lázari et al. 2021. “LncRNAs of *Saccharomyces cerevisiae* dodge the cell cycle arrest imposed by the ethanol stress”. BioArxiv. https://doi.org/10.1101/2021.06.28.450142.

